# STADiffuser: high-fidelity simulation and full-view 3D modeling of spatial transcriptomics

**DOI:** 10.1101/2025.08.30.673245

**Authors:** Chihao Zhang, Shihua Zhang

## Abstract

Spatial transcriptomics technologies have provided invaluable insights by profiling gene expression alongside precise spatial information. However, they encounter high costs, data sparsity, and limited resolution, hindering their broader adoption and utility. Moreover, the lack of flexible simulators capable of generating high-fidelity simulated data has impeded the development of computational tools for spatial transcriptomic data analysis. To this end, we introduce STADiffuser, a versatile deep generative model that leverages diffusion modeling for accurate simulation of spatial transcriptomic data.

STADiffuser employs a two-stage architecture: an autoencoder with a graph attention mechanism for learning spot embeddings, followed by a latent diffusion model integrated with a spatial denoising network for data generation. STADiffuser is the first simulator designed for spatial transcriptomic data, capable of handling multiple samples and 3D coordinates while supporting user-defined conditions. STADiffuser facilitates various downstream analyses, including accurate imputation, super-resolution, and full-slice generation.

Furthermore, its generative scheme enables *in silico* experiments, thereby enhancing the statistical power in detecting differentially expressed genes and identifying cell-type-specific genes while effectively controlling confounding factors. Notably, STADiffuser scales to millions of spots and supports the full-view 3D modeling of a marmoset cerebellum atlas with over 20 million spots, enabling detailed investigations from arbitrary viewing angles.

## Introduction

Spatial transcriptomics (ST) technologies can now profile gene expression from hundreds to thousands of genes in parallel alongside high-resolution spatial information. For instance, 10x Visium^1^ achieves a spatial resolution of 55 µm, with each captured location (referred to as spot in the following) referring to 1-10 cells. Advancements in barcoding-based methods, such as Slide-seq^2,3^, which achieves spatial resolutions approaching cellular levels (10 µm), and Stereo-seq^4^, which achieves subcellular resolutions (0.22 µm), have significantly improved spatial resolution capabilities. In addition, Slide-tags^5^, a recent innovative technique, has accomplished single-nucleus resolution-level profiling of gene expression and epigenetic regulation within individual cells.

The advancements of those ST technologies have revolutionized biology investigation and motivated the development of numerous computational methods to facilitate in-depth analysis. Those newly developed methods can identify spatial regions with similar expression profiles^6,7^, detect genes exhibiting spatial variability^8–10^, characterize cell-cell communications, conduct multi-slice analysis for analyzing 3D gene expression patterns^11,12^, and enhance the spatial resolution of the ST data^13,14^.

However, several inherent challenges persist. First, ST technologies incur considerable costs, posing a significant economic burden for laboratories and hindering their widespread adoption and application. Additionally, ST data suffers from excessive sparsity and inadequate resolution. Due to technical resolution, ST technologies struggle to match the resolution of single-cell mRNA sequencing (scRNA-seq)^15^, posing significant challenges for subsequent biological analysis. Furthermore, the development of computing tools also urgently requires a flexible simulator to facilitate benchmarking.

Consequently, generating high-fidelity simulated ST data emerges as a critical problem from a computational perspective. Such simulators can help develop computing tools, provide information for experimental design, and effectively reduce experimental costs. Moreover, they also have the potential to enhance ST data by addressing its inherent sparsity and limited resolution. Most importantly, generative models could be powerful tools for analyzing spatial transcriptomics data, uncovering underlying patterns, and facilitating *in silico* experiments.

While simulating realistic transcriptomics has been studied in scRNA-seq data^15–18^, spatial transcriptomics introduces unique challenges that limit the applicability of existing simulators. The direct application of these tools often leads to a loss of spatial structure^19^. Although some simulators attempt to incorporate spatial information—for instance, ZINB-WaVe^16^ models spatial coordinates as covariates in a zero-inflated negative binomial framework, and scDesign3^20^ employs copula models to simulate single-cell and spatial omics data jointly—they still rely on parametric assumptions. They cannot tailor to the unique requirements of ST data. SRTSim^19^, perhaps the first ST-specific simulator, generates high-fidelity simulated data by reordering simulated count data based on expression level ranks to preserve spatial patterns.

However, existing simulation methods—whether parametric or non-parametric—share several intrinsic limitations. Parametric approaches, such as ZINB-WaVe and scDesign3, rely on strong distributional assumptions (e.g., Poisson or negative binomial), which may be violated in complex real-world ST data. While non-parametric methods like SRTsim avoid such assumptions, they still exhibit limited model capacity, making it hard to learn rich spatial dependencies, adapt to diverse tissue architectures, or generalize across multiple samples and 3D spatial contexts. Moreover, most existing methods are not designed to scale efficiently to datasets with millions of spatial spots, constraining their applicability in large-scale simulations and full-view spatial atlas reconstruction.

To fill those gaps, we present STADiffuser, a deep generative model enabling high-fidelity ST data generation while incorporating multiple samples and 3D coordinates within a unified framework. STADiffuser is the first ST-specific simulator capable of multi-slice and 3D modeling. Extensive experiments demonstrate that it could generate realistic data with user-specified conditions and empower various downstream analyses. Importantly, STADiffuser scales to millions of spatial spots, making it suitable for large-scale simulations and the reconstruction of a full-view 3D atlas.

## Results

### Overview of STADiffuser

STADiffuser is a flexible deep generative model designed to generate high-fidelity ST data by fully utilizing spatial information. It comprises two key components: a graph attention autoencoder, which learns the embeddings of the spots, and a spatial denoising network that facilitates denoising in the latent embedding space (**Fig. 1a**). The training process of STADiffuser occurs in two stages. Initially, it constructs a spatial neighbor network (SNN) using the spots’ coordinates to capture local spatial information. Subsequently, a deep graph attention autoencoder is employed to learn spot embeddings. For multi-slices, we introduce a triplet learning strategy to train the autoencoder. This approach can mitigate the batch effect and align the spot embeddings across sections. In the second stage, STADiffuser adopts a latent space diffusion model, which gradually transforms random Gaussian noise into the distribution of spots in the latent space. STADiffuser proposes a spatial denoising network that integrates the coordinates of spots through spatial encoding with optional conditioning labels such as cell type or domain labels to enhance the transformation process.

**Fig. 1.**
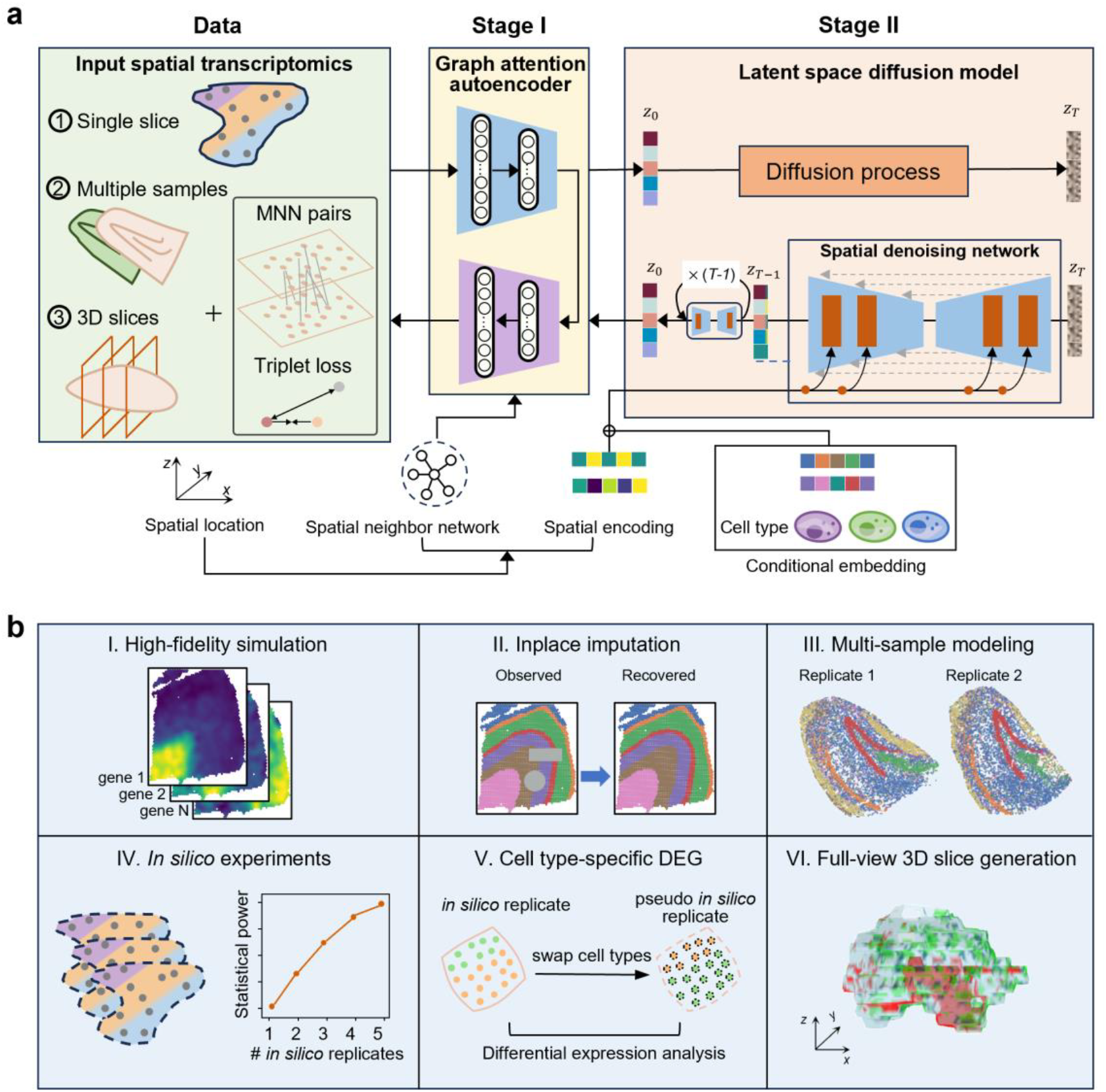
Overview of STADiffuser. **a**, STADiffuser is a two-stage procedure tailored specifically for ST data modeling. In the first stage, it learns spot embeddings using an autoencoder without presuming the data distribution. This autoencoder is enhanced with a graph attention mechanism^21^, enabling it to adaptively aggregate information of neighboring spots when needed. In the second stage, a latent diffusion model^22,23^, coupled with a spatial denoising network for the spot embedding space, is trained. The spatial denoising network learns to progressively transform the standard Gaussian into the distribution of spot embedding space. 2D/3D spatial coordinates are encoded and integrated into the denoising network to preserve long-range spatial dependencies. Finally, simulated data are generated by decoding the simulated embeddings into the gene expression space. STADiffuser is scalable, supporting various configurations, including ➀ single slices, ➁ multiple samples, and ➂ 3D slices. **b**, STADiffuser is a versatile generative model that empowers high-fidelity spatial transcriptomic data simulation, in-place gene spatial profile imputation, multi-sample modeling, *in silico* experiments, cell-type specific differential gene analysis, and full-view 3D *de novo* generation.

During the sampling phase, STADiffuser first generates Gaussian noise. Its trained denoising network progressively transforms this Gaussian noise, alongside specified spatial coordinates and conditioning labels, into the distribution of latent spot embeddings. In the final step, *in silico* ST data is obtained by decoding the simulated latent embeddings with the trained decoder. STADiffuser empowers various downstream analyses and facilitates in-depth biological investigations (**Fig. 1b**).

### STADiffuser enables high-fidelity generation and imputation within the human dorsolateral prefrontal cortex

To assess the effectiveness of STADiffuser, we first applied it to a human dorsolateral prefrontal cortex (DLPFC) dataset, which was profiled using the 10x Visium platform^24^ and manually annotated based on marker genes and morphological images. The human DLPFC is layer-structured with six layers and white matter (WM). We illustrate one section (151767) consisting of 3,460 spots (**Fig. 2a**). We kept the top 3000 highly variable genes and evaluated the performance of STADiffuser and five baseline methods (i.e., Splatter, KerSplatter, ZINBwave, scDesign3, and SRTSim). Among these methods, Splatter and KerSplatter were designed for scRNA-seq simulation without incorporating spatial information. ZINBwave and scDesign3 are applicable for both scRNA-seq and ST data. SRTSim is a nonparametric statistical approach tailored for ST simulation. We computed the gene-wise Pearson correlation between the simulated data and the enhanced raw data to measure the fidelity of how closely they are.

**Fig. 2.**
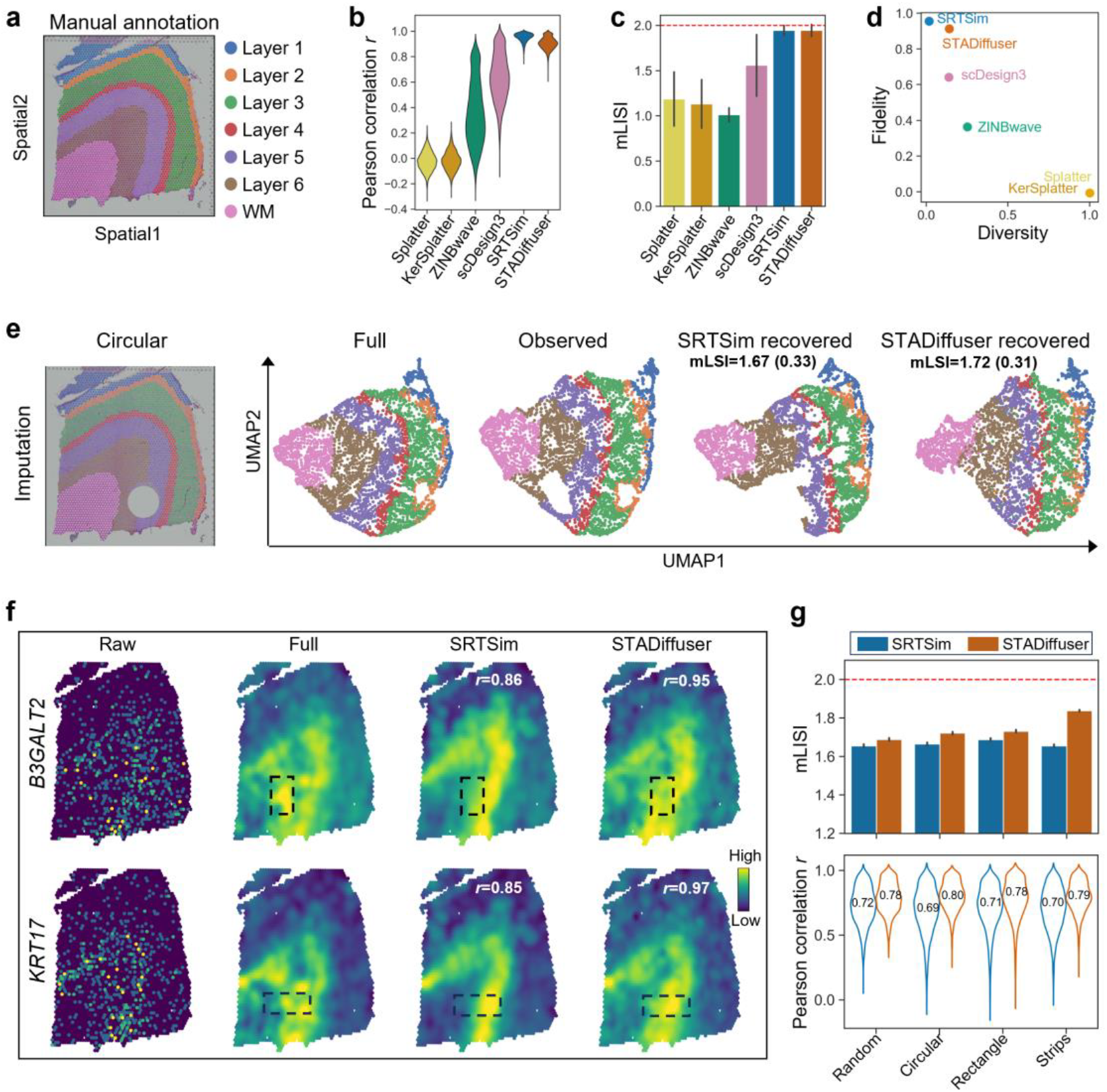
STADiffuser enables high-fidelity generation and imputation within the human dorsolateral prefrontal cortex dataset. **a**, Manual annotation of cortical layers and white matter (WM) in section 151676 of the DLPFC dataset. **b**, Violin plot illustrating Pearson correlation between the real data and the data generated by STADiffuser along with five baseline methods. **c**, Bar plot showing the mean of local inverse Simpson Index (mLISI) of STADiffuser and the five methods, with the optimal mLISI=2 indicated by a horizontal dashed line. **d**, Scatter plot depicting the characteristics of simulation methods in terms of diversity (variation among simulated data) and fidelity (similarity of simulated data to real data). **e**, Comparison of imputation performance on a section with a masked hole, including the spatial mapping and the UMAP embeddings of the full, observed, and recovered data. **f**, Expression profiles of representative genes across raw, full, and recovered data with Pearson correlation annotated. **g**, Comparative analysis of SRTsim and STADiffuser imputation performance across various scenarios.

We also calculated the mean of spot-wise Local Inverse Simpson’s Index (mLISI) to assess how well the simulated data preserve the latent topology in embedding space. Notably, a desired simulator should generate data that resembles raw data and maintains diversity, indicating the variations and heterogeneity inherent in biological samples. To measure the diversity of simulators, we defined the diversity metric as 1 − *r*, where *r* is the pairwise Pearson correlation of the simulated replicates. Please refer to the subsection “**Evaluation metrics of simulation methods**” for a detailed description of the evaluation metrics.

Among the compared methods, SRTSim and STADiffuser outperform the others in terms of gene-wise Pearson correlation (**Fig. 2b**). Consistently, SRTSim and STADiffuser also effectively preserve the latent topology, as indicated by their mLISI values approaching the optimal value of 2 (**Fig. 2c**). Methods like Splatter and KerSplatter, which disregard spatial information, fail to generate simulated data that resemble the actual data, highlighting the importance of spatial information in data fidelity. It is worth noting that there is a trade-off between fidelity and diversity. As shown in **Fig. 2d**, methods of high diversity typically result in low fidelity. STADiffuser outperforms SRTSim in terms of diversity while remaining high fidelity.

Although nonparametric statistical approaches such as SRTSim demonstrate satisfactory fidelity, we posit that deep generative methods hold more promise in tackling challenging scenarios due to their capacity to learn the underlying distribution. Hence, we evaluated the performance in imputing the missing regions. We removed a circular area from the section (**Fig. 2e**, left panel) and applied SRTSim and STADiffuser to this section to generate the missing spots by specifying their original coordinates. STADiffuser accurately recovered the UMAP embeddings of the observed data, making them closely resemble those of the full data, achieving an mLISI value of 1.72 (**Fig. 2e**).

However, SRTSim failed to fill the missing hole in the observed UMAP embeddings. STADiffuser also outperforms SRTSim in terms of reconstructing the spatial distribution of representative genes, including *B3GALT2* and *KRT17* (**Fig. 2f**). We evaluated the imputation performance under different scenarios, including random missing, circular, rectangle and strips (**Fig. 2g**).

STADiffuser consistently outperforms SRTSim in terms of latent topology preservation (**Fig. 2g**, upper panel) and gene imputation (**Fig. 2g**, lower panel). We further investigated the performance of the two methods with varying random missing rates (**Fig. S1**), and the results were still consistent. In short, our findings highlight the potential of deep generative methods over nonparametric approaches such as SRTSim, particularly in addressing challenging scenarios. Please refer to the **Supplementary Materials** for a detailed description of the experimental settings.

### STADiffuser facilitates multi-sample modeling and enhances analysis of the hippocampus in mouse models of Alzheimer’s disease

To showcase the proficiency of STADiffuser for multi-sample modeling, we applied it to a mouse hippocampus dataset obtained from the J20 mouse model of Alzheimer’s disease (AD)^25^. Four biological replicates of the J20 mouse model profiled by Slide-seq V2 were manually annotated with marker genes (**Fig. 3a** and **Fig. S2a**). The number of spots detected in each replicate ranges from 10,000 to 16,000. We identified 3,000 top highly variable genes in each replicate and determined their intersection. Consequently, 2,455 genes were shared across all replicates. The spatial coordinates of the four replicates are not aligned, and the anatomical structure varies in shape. We manually aligned the spatial coordinates of replicates 1 and 2 by referencing the distinct `V’ shape of the dentate gyrus (DG, colored in red). Replicates 3 and 4, however, exhibited significant morphological variation and could not be aligned by simple geometric transformation. Thus, they were excluded from alignment. As a result, 23,916 spots are remaining. STADiffuser can enable the embeddings of the two replicates to be well aligned and mixed (**Materials and Methods, Fig. S2c**). We simulated an *in silico* replicate, denoted as `STADiffuser (ms)’ of replicate 1, by specifying the original coordinates and cell type. In parallel, we trained an integrative representation model across all four biological replicates using an autoencoder with a triplet loss objective.

**Fig. 3.**
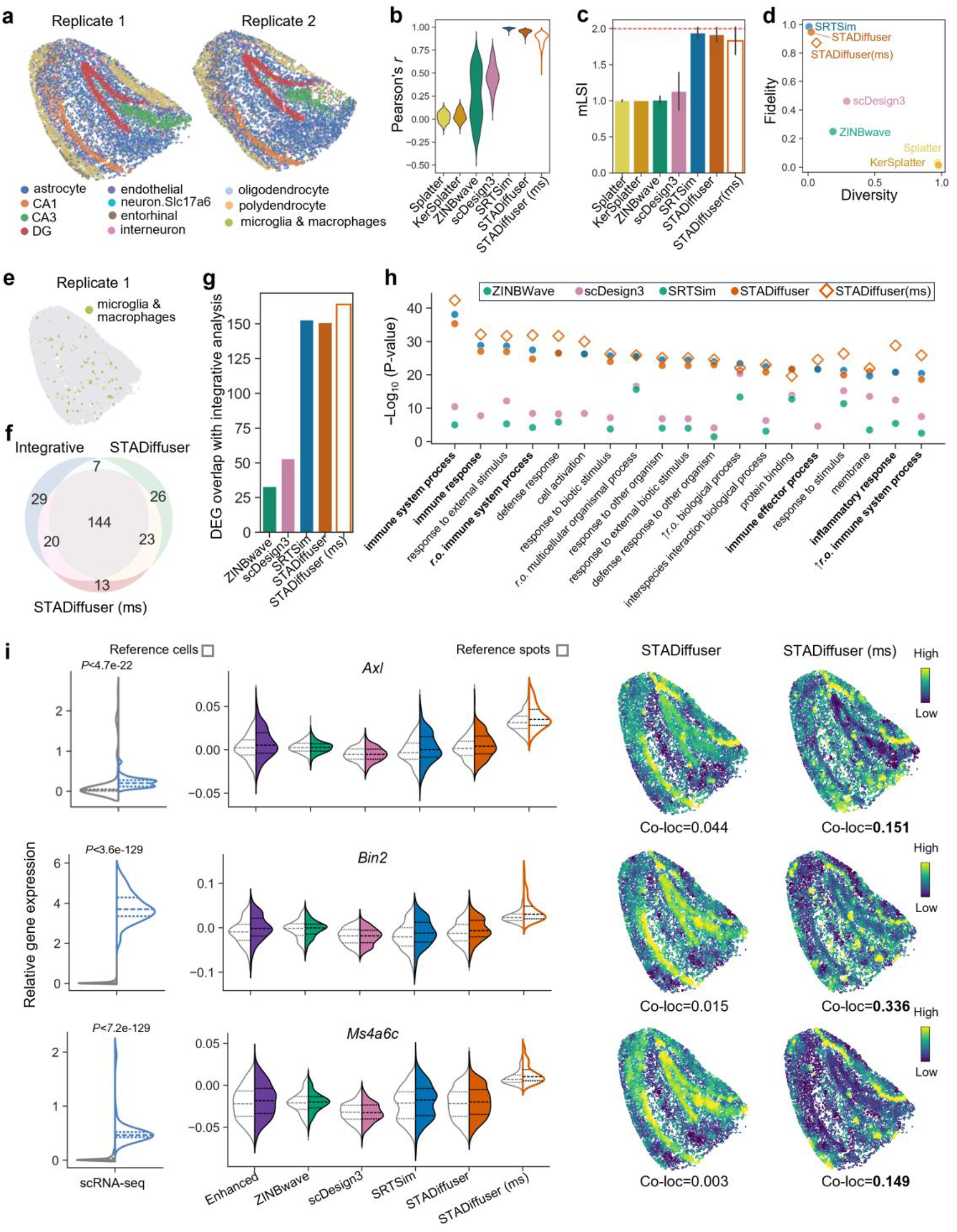
STADiffuser facilitates multi-sample modeling and enables in-depth analysis of the hippocampus in mouse models of Alzheimer’s disease. **a**, Spatial mapping of the two biological replicates of the J20 mouse’s hippocampus. **b**, Violin plot illustrating the Pearson correlation between the real data and the data generated by STADiffuser, along with five baseline methods. **c**, Bar plot showing the mLISI of the simulation methods, with the optimal mLISI=2 indicated by a horizontal dashed line. **d**, Scatter plot illustrating the characteristics of simulation methods; multi-sample (ms) STADiffuser, indicated with a diamond marker, showing increased variation and decreased fidelity. **e**, Spatial mapping of microglia and macrophage cells on replicate 1. **f**, Bar plot showing the overlap of differentially expressed genes (DEGs) detected in simulated data with DEGs identified from integrative analysis based on reconstructed expression from four real biological replicates. **g**, Venn diagram illustrating the overlap of DEGs among STAdiffuser, STAdiffuser (ms), and integrative analysis. **h**, Dot plot displaying the top GO enrichment analysis of DEGs on simulated data, indicating that DEGs from STAdiffuser (ms) are more enriched in immune-related terms. **i**, Violin plots comparing the expression profiles of three representative genes in both the scRNA-seq reference data and the simulated data for microglia and macrophage cells/spots with reference cells/spots. Co-localization values, indicating the spatial consistency between microglia and macrophage spots and the simulated highly expressed genes, are annotated in the spatial maps in the last two columns.

This model served as the basis for downstream analyses, including differential gene expression. As the comparison methods lack support for multi-sample modeling, we restricted their application to replicate 1.

We first evaluated the impact of using multiple samples during simulation. As expected, using multiple biological replicates increases the variability in the simulated data, but it also slightly reduces the fidelity (**Fig. 3b-d**). This trade-off is reasonable, as biological variability introduces additional complexity.

Nevertheless, STADiffuser with multi-sample input still ranked closely behind STADiffuser and SRTSim, while maintaining a clear fidelity advantage over other baseline methods (**Fig. 3d**).

Microglia and macrophage cells are critical in the progression of Alzheimer’s disease^26^. In our study, we conducted a case study to identify differentially expressed genes (DEGs) for them (**Fig. 3e**). Initially, we validated that integrative analysis performed on the reconstructed expression profiles— obtained by training an autoencoder with triplet loss across four replicates— yielded more significant DEGs compared to using a single replicate alone (**Fig. S2d**). Moreover, integrated analysis led to the identified DEGs being more biologically relevant, as evidenced by significant enrichment in immune-related and microglia-related Gene Ontology (GO) terms (**Fig. S2e**). Using the integrative DEGs as a reference, we observed that STADiffuser with multiple samples exhibited a distinct overlap compared to when only one replicate was used (**Fig. 3f**), as well as compared to other methods (**Fig. 3g**). Furthermore, we conducted gene enrichment analysis for the DEGs identified within the simulated data. As illustrated in **Fig. 3h**, DEGs identified by STADiffuser (ms) showed significantly higher enrichment in immune-related GO terms than other methods.

We reasoned that the simulated data using multiple samples would be more biologically plausible. To validate this, we obtained a J20 mouse model hippocampus single-cell RNA-seq dataset from another study^27^. This dataset comprised 2,933 cells, and we retained the same gene set as in the previous analysis. We employed the top 200 marker genes identified in microglia and macrophages to score the cells. We successfully identified the corresponding cell clusters in the scRNA-seq dataset, as depicted in the UMAP embeddings (**Fig. S2b**, marked with red circles). As the single-cell sequencing dataset provides higher resolution by capturing gene expression profiles at the level of individual cells, with it as a reference, we plotted the gene expression levels in microglia and macrophage cells against the rest (**Fig. 3i**, left panels). Specifically, the *Axl* gene plays a crucial role in microglia and macrophage cells and modulates inflammatory responses^26^. While single-cell RNA sequencing data demonstrates its high expression, the limited sensitivity of spatial transcriptomic sequencing platforms resulted in capturing only a few reads of Axl expression. Consequently, neither the raw data enhanced by the autoencoder trained on a single replicate nor the simulated data using only one replicate identified *Axl* as a DEG. However, in the simulated data generated by STADiffuser with multiple samples, *Axl* was significantly more highly expressed than in the reference spots (**Fig. 3i**, first row, middle panel). Similarly, *Bin2*, a member of the TYROBP signaling pathway dysregulated in microglia during Alzheimer’s^28^, was identified only using scRNA-seq data or STADiffuser (ms) simulated data (**Fig. 3i**, second row). The same observation applies to *Ms4a6c*, a member of the *Ms4* gene family that plays a role in Alzheimer’s disease risk, potentially affecting neuroinflammation and β-amyloid clearance through monocytes and microglia^29^. Moreover, genes stimulated by STADiffuser (ms) exhibit more distinct spatial enrichment around microglia and macrophage cells. While utilizing only a single replicate fails to capture these spatial patterns (**Fig. 3i**, right panels). To quantify spatial consistency between the simulated genes and microglia & macrophage cells, we introduced a ‘co-localization’ (co-loc) metric (see subsection “**Computing the co-localization metric**”). It is evident that the spatial consistency of genes simulated by STADiffuser (ms) with microglia and macrophage cells is significantly higher compared to other methods in most cases (**Fig. S3b**). Moreover, most of the 30 unique DEGs identified in STADiffuser (ms) simulated data showed significant differential expression in scRNA-seq data, affirming the biological plausibility of employing multiple replicates in simulations (**Fig. S3c-d**). In summary, simulating data with multiple biological replicates increases variability and enhances statistical power, helping to identify biologically relevant genes that may be missed with a single replicate.

### STADiffuser-generated *in silico* replicates enhance biological discovery in the mouse visual cortex

We applied STADiffuser to a mouse visual cortex dataset generated by STARMap^30^, an *in situ* imaging-based spatial transcriptomic technology, to demonstrate its power in enhancing biological analysis. This dataset comprises 1,020 genes and 1,207 cells. The section was manually annotated (**Fig. 4a**). In terms of gene-wise Pearson correlation, STADiffuser simulated data ranked second only to SRTSim (**Fig. 4b**) and slightly outperformed SRTSim in preserving latent space topology (**Fig. 4c**). Notably, STADiffuser occupies a unique position by demonstrating much higher diversity in generated data compared to SRTSim, while maintaining high fidelity, ranking second only to SRTSim (**Fig. 4d**).

**Fig. 4.**
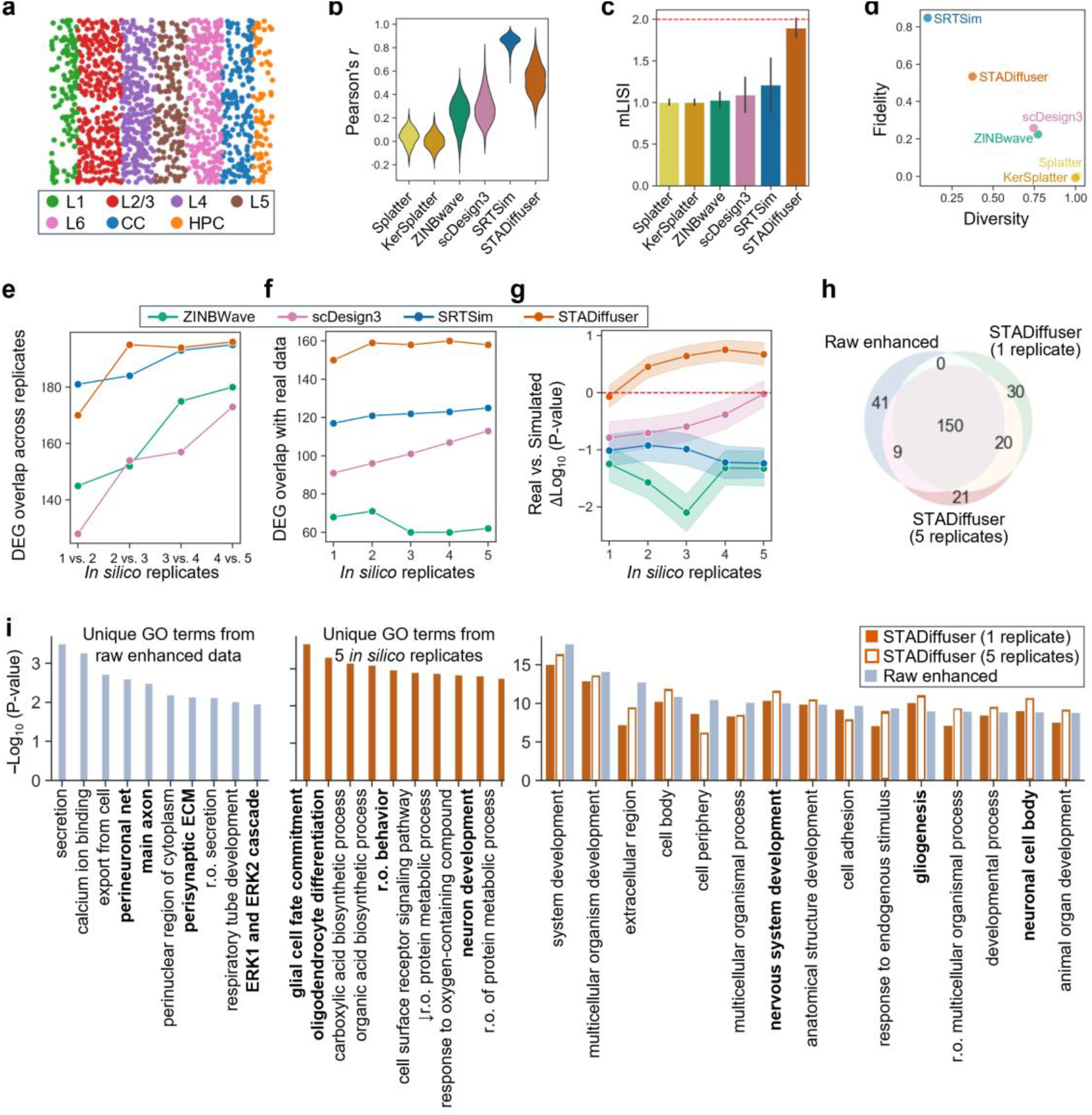
STADiffuser-generated *in silico* replicates enhance biological discovery in the mouse visual cortex dataset. **a**, Spatial mapping of the mouse visual cortex. **b**, Violin plot illustrating the Pearson correlation between the real data and the data generated by simulation methods. **c**, Bar plot showing the mLISI of the simulation methods, with the optimal mLISI indicated by a horizontal dashed line. **d**, Scatter plot illustrating the characteristics of simulation methods. **e**, Overlap of top 200 DEGs across successive *in silico* replicates. **f**, Overlap of top 200 DEGs between raw enhanced data and increasing *in silico* replicates. **g**, Differences in log^10^(*P*-value) of enriched GO terms between the real and simulated data concerning increasing *in silico* replicate numbers; the 95% confidence intervals are represented by shading. **h**, Venn diagram illustrating the overlapping number of DEGs between raw enhanced data and STADiffuser simulated data. **i**, Bar plot displaying the −log10(*P*-value) of the top ten unique GO terms from raw enhanced data and STADiffuser simulated data (five replicates), as well as the top 15 commonly enriched GO terms.

To examine the impact of the increase in the number of *in silico* replicates, we analyzed DEGs in the hippocampus (HPC) layer with simulated replicates ranging from 1 to 5. Specifically, the overlaps of the top 200 DEGs across consecutive increasing *in silico* replicates tend to increase (**Fig. 4e**), implying that using multiple replicates enhances the reliability and robustness of DEG analysis. Using the top 200 DEGs identified in raw data enhanced by autoencoder as a reference, we calculated the overlap of the top 200 DEGs identified within simulated data (**Fig. 4f**). All the methods excluding ZINBWave display an increased trend in the overlap with the raw data reference. In addition, despite data generated by SRTSim having a higher Pearson correlation than STADiffuser, the DEG overlap of STADiffuser simulated data is significantly higher than that of SRTSim. Arguably, STADiffuser is more suitable for downstream biological analysis tasks than SRTSim.

We further calculated the differences in the logarithmic *P* values between the raw and simulated data. It turns out that the significance levels of GO terms enriched in DEGs identified from STADiffuser-simulated data exceed those from the raw enhanced data, particularly when more than one replicate is used (**Fig. 4g**). This suggested that STADiffuser may capture the biological variation, thereby refining the results of DEG analysis. In addition, the DEGs identified from five STADiffuser *in silico* replicates show greater overlap with the reference than those from a single replicate (**Fig. 4h**). In contrast, none of the compared methods resulted in significantly higher P-values of enriched GO terms than those of raw enhanced data, even when varying the number of replicates from 1 to 5 (**Fig. S4**). Specifically, SRTSim exhibited a decrease in significance levels of enriched GO terms, consistently falling below those of the raw enhanced data. Meanwhile, the significance levels of GO terms in scDesign3-simulated data increased to match those of the raw enhanced data. As a result, although scDesign3 has a higher variation than STADiffuser, it failed to improve biological discovery.

We visualized the GO terms enriched in DEGs of the raw enhanced data, as well as from STADiffuser-simulated data using one and five *in silico* replicates, to evaluate how increasing the number of replicates influences both the biological relevance and enrichment significance of discovered GO terms. GO terms, such as glial cell fate commitment and oligodendrocyte differentiation, that are uniquely enriched in DEGs of five STADiffuser replicates are more biologically relevant to HPC (**Fig. 4i**, middle panel, specifically relevant terms shown in bold). In contrast, GO terms uniquely enriched in raw enhanced data comprise more general terms that are not specifically related to HPC, including secretion and calcium ion binding.

Compared to using a single replicate, DEGs identified by STADiffuser within five replicates were more significantly enriched in almost all GO terms (**Fig. 4i**, right panel). Furthermore, among the top 15 GO terms enriched in both STADiffuser-simulated and raw enhanced data, terms specifically related to HPC, such as nervous system development, gliogenesis, and neuronal cell body, were enriched at more significant levels than in the raw enhanced data (**Fig. 4i**, right panel; specifically relevant terms shown in bold). Hence, STADiffuser potentially captures biological variation, and its use of multiple *in silico* replicates may enhance the power of DEG detection, thereby improving biological discovery through *in silico* simulation.

### STADiffuser empowers the identification of cell-type-specific genes and unveils the human melanoma tumor heterogeneity in single-nucleus spatial profiles

Identifying cell-type-specific marker genes is crucial for delineating tissue sections in tumor heterogeneity. However, this task is challenging due to the complex interplay between gene expression, cell types, spatial localizations, and other confounding factors. STADiffuser presents an intuitive method for pinpointing cell-type-specific genes while effectively addressing localization confounders, providing a deeper insight into tumor heterogeneity. Here, we used a human melanoma tumor dataset obtained through Slide-tags profiling with 6,466 cells. We retained the top 2,000 highly variable genes. Notably, the tissue section exhibited a high degree of cellular heterogeneity with multiple cell types spatially intermixed. The UMAP embeddings of gene expression revealed distinct clusters (**Fig. 5a**) courtesy of Slide-tags’ single-nuclei resolution. The observations remain consistent with previous experiments.

**Fig. 5.**
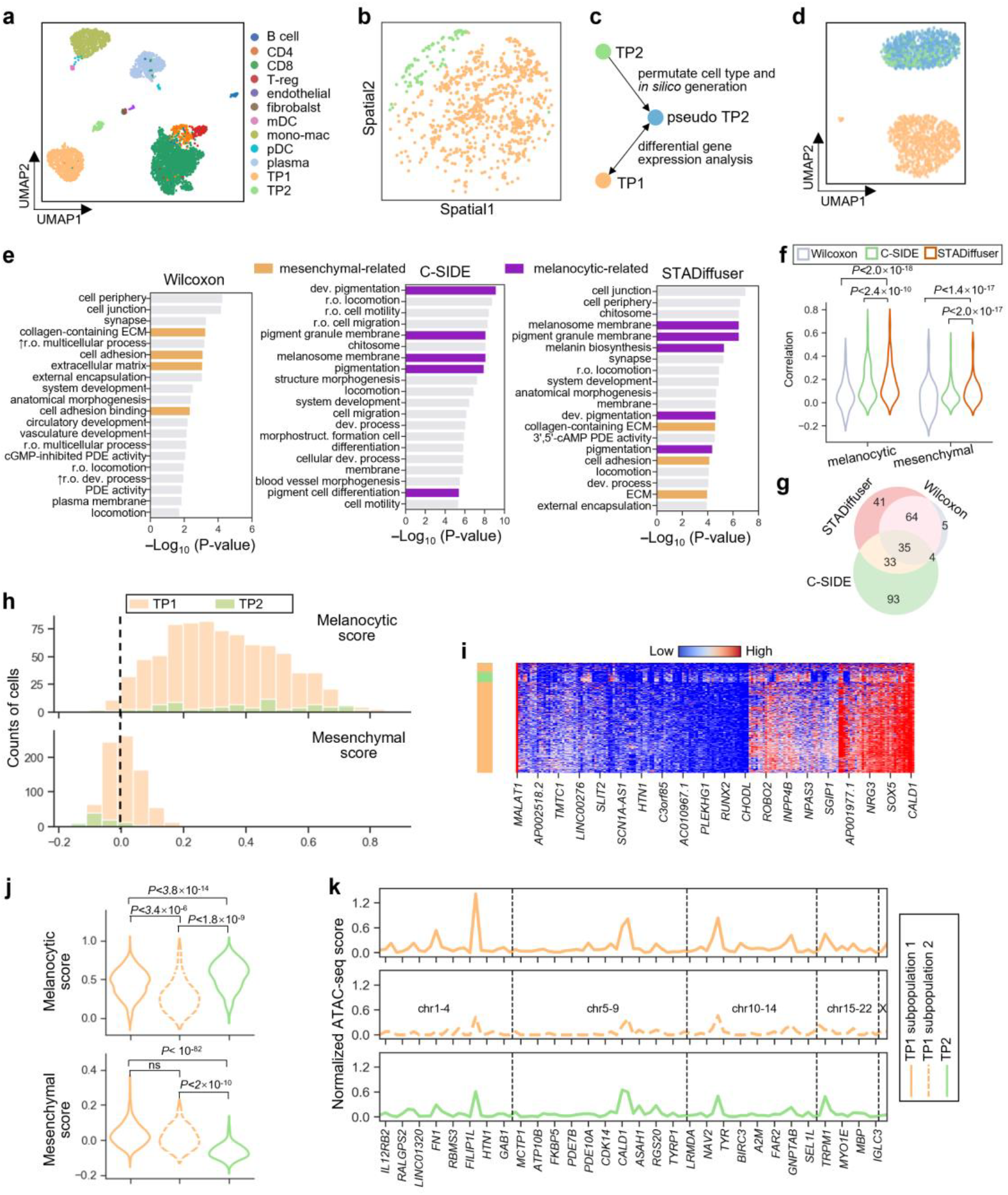
STADiffuser empowers the identification of cell-type-specific DEGs and unveils the heterogeneity of human melanoma tumors in single-nucleus spatial profiles. **a**, UMAP embeddings of the snRNA-seq profiles colored by cell types. **b**, Spatial mapping of TP1 and TP2. **c**, Schematic overview of the STADiffuser pipeline for cell-type-specific DEG identification. **d**, UMAP embeddings of TP1, TP2, and the generated pseudo-TP2, colored as in panel **c. e**, Bar plots presenting the top 20 enriched GO terms identified by the Wilcoxon test, C-SIDE, and STADiffuser. STADiffuser uniquely enriches both mesenchymal-related and melanocytic-related terms. **f**, Violin plots depicting correlations between DEGs identified by three different methods and both melanocytic and mesenchymal scores. **g**, Venn diagram illustrating the overlap of the DEGs identified by the Wilcoxon test, C-SIDE, and STADiffuser. **h**, Histograms depicting the distribution of melanocytic and mesenchymal scores across spots of TP1 and TP2. **i**, Heatmap illustrating the gene profiles of cell-type-specific DEGs identified by STADiffuser; TP1 displays approximately two clustered subpopulations based on the identified DEGs. **j**, Violin plots showing the distribution of melanocytic and mesenchymal scores in subpopulations 1 and 2 of TP1 and TP2. **k**, Normalized chromatin accessibility across STADiffuser-identified cell-type-specific genes in subpopulations 1 and 2 of TP1 and TP2 overall; the chromatin regions are delineated by dashed vertical lines.

Among them, the simulated data by STADiffuser exhibited notably high diversity and ranked second only to SRTSim in reconstructing gene correlation (**Fig. S5a-c**).

We observed two distinct tumor populations, namely TP1 and TP2 (**Fig.5b**), with TP1 comprising significantly more cells than TP2. To mitigate the potential confounding effect of cell-type localization, we applied STADiffuser to generate pseudo-TP2 cells by swapping the cell type assignments between TP1 and TP2 (**Fig. 5c**). This *in silico* approach produced pseudo-TP2 replicates with cell counts comparable to TP1. UMAP embeddings showed that TP1 and TP2 are distinct, while TP2 and pseudo-TP2 are well mixed (**Fig. 5d**). Using those data, we then identified 172 DEGs by comparing gene expression levels between TP1 and pseudo-TP2 cells. For comparison, the Wilcoxon rank-sum test (which does not consider localizations) and C-SIDE^25^ (a statistical approach to accounting for localizations by a generalized linear model) identified the same number of DEGs.

Prior analysis^5^ suggest that tumor cells fall into two main cell states: melanocytic-like and mesenchymal-like. Interestingly, the Wilcoxon rank-sum identified genes were primarily enriched in mesenchymal-related GO terms, while C-SIDE identified genes were mainly enriched in melanocytic-related GO terms.

In contrast, STADiffuser-identified genes were uniquely enriched in GO terms associated with both cell states. Those genes exhibited greater overlap with those identified by the Wilcoxon rank-sum test. While a significant portion of C-SIDE-identified genes were exclusive to it (**Fig. 5g**). To further evaluate biological relevance, we scored TP1 and TP2 cells using melanocytic and mesenchymal marker genes adapted from previous works^31,32^. We calculated the correlation between the identified gene expression levels and those scores. STADiffuser-identified genes showed significantly stronger correlations with both cell-state scores, suggesting that they more accurately capture the distinct transcriptional programs of the two tumor populations (**Fig. 5f** and **Fig. S5f)**.

The histograms of melanocytic-like and mesenchymal-like scores depicted distinct profiles for the two tumor cell populations **(Fig. 5h**): TP2 cells were primarily melanocytic-like, whereas TP1 cells comprised both melanocytic-like and mesenchymal-like cells. This observation was supported by GO enrichment analysis. Notably, the Wilcoxon rank-sum test, when not accounting for localizations, primarily identified genes associated with the mesenchymal-like state (**Fig. 5e**, left panel). While C-SIDE predominantly identified genes enriched in melanocytic-related GO terms, seemingly overcorrecting the localization effect. Furthermore, STADiffuser identified genes that clustered the TP1 cell into approximately two subpopulations (**Fig. 5i**), unlike the other two methods, which failed to distinguish these subpopulations (**Fig. S5e**). The two identified subpopulations exhibited marked differences in melanocytic scores (**Fig. 5j**, upper panel). We analyzed another adjacent section using the multiome Slide-tags technology, which simultaneously measures ATAC-seq and RNA-seq, to validate our findings.

The spatial distribution of the two tumor cell populations, TP1 and TP2, revealed that TP1 consistently clusters into two subpopulations based on genes previously identified by STADiffuser (**Fig. S5h, i**). The cell type distributions between the two subpopulations show a distinct difference, as indicated by the fold change in cell typing co-localization (**Fig. S5g**).

Moreover, the ATAC-seq profiles also revealed two distinct epigenetic differences among the TP1 subpopulations and TP2 cells (**Fig. 5k**).

To conclude, STADiffuser provides a powerful tool for pinpointing cell-type specific genes that advance our understanding of tumor heterogeneity. We argue that the localization effect could be complicated and nonlinear.

Therefore, in contrast to the statistical approach relying on generalized linear models, such as C-SIDE, the deep generative approach shows promise in capturing these nonlinear effects more effectively, thereby identifying signature genes that define tumor heterogeneity.

### STADiffuser performs accurate whole-slice 3D imputation in the *Drosophila* embryo

The advent of high-resolution ST technologies, such as Stereo-seq, has enabled the creation of 3D spatial maps of complex biological systems. Nevertheless, current simulation methods face constraints in accurately capturing these 3D slices. A notable strength of STADiffuser is its ability to model 3D slices within a unified framework, offering a more reliable approach to data generation and reconstruction.

To showcase its ability, we used a Drosophila embryo dataset (16-18 h after egg laying, termed E16-18) profiled by Stereo-seq^33^. This dataset contains 14 cryosection slices, 7 μm thick, with manually annotated cell types (**Fig. 6a**). Each slice contains approximately 1,000 spots, totaling 14,634 overall. Initially, we retained the top 2,000 highly variable genes and trained an autoencoder with triplet loss to mitigate the batch effect across slices. We trained a latent diffusion model with a spatial denoising network with 3D positional embedding. The resulting low-dimensional embeddings showed clear separation of cell types within the aligned UMAP space (**Fig. 6b**). The well-mixed UMAP visualization of raw and simulated data demonstrates high simulation fidelity with an mLISI value of 1.88 (an optimal value of 2 indicates perfect mixing of raw and simulated data).

**Fig. 6.**
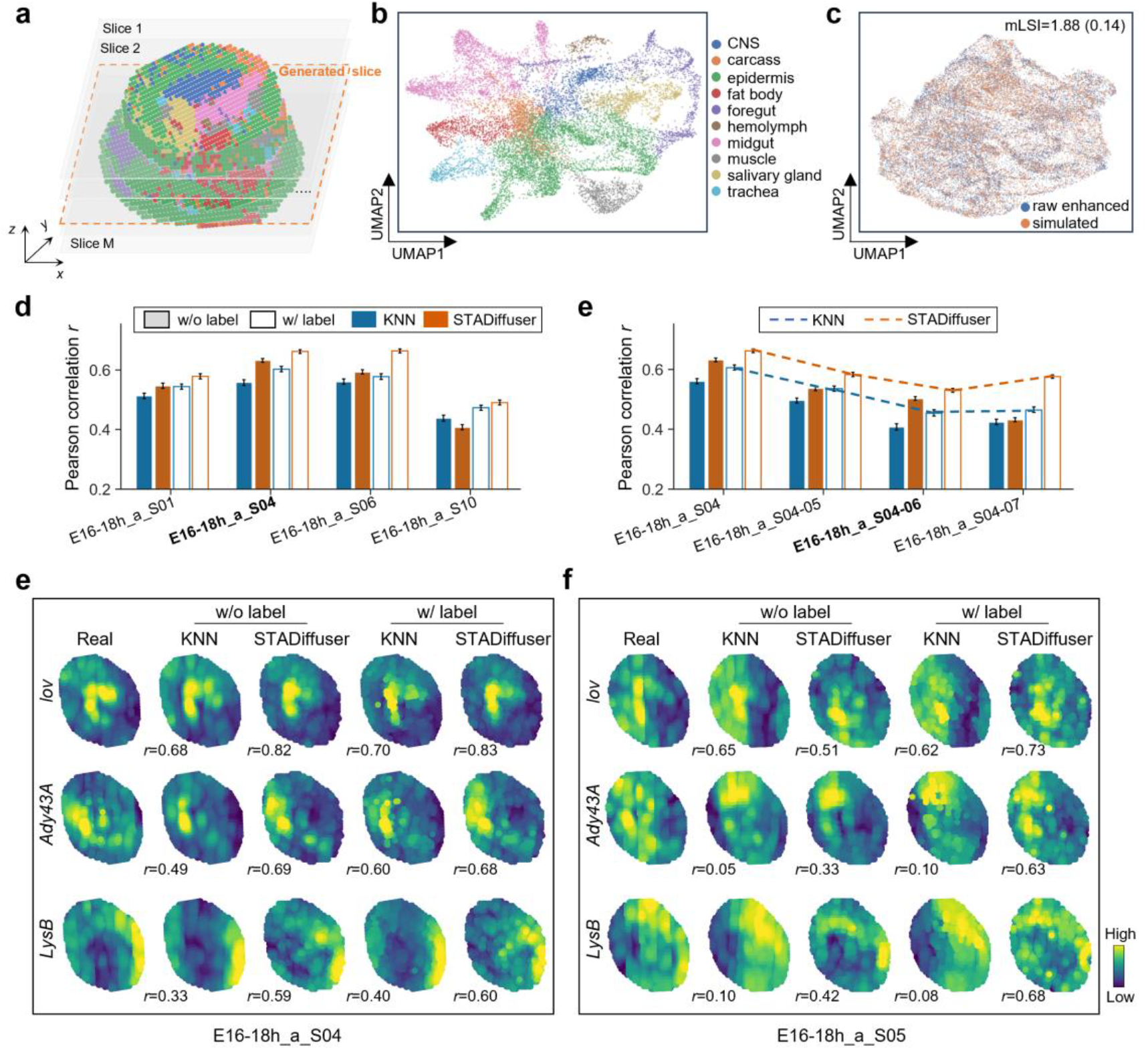
STADiffuser facilitates 3D slice generation in the *Drosophila* embryos dataset. **a**, 3D spatial mapping of E16-18h *Drosophila* embryos, colored by cell types. The generated slice is highlighted with an orange border. **b**, UMAP embedding of aligned spot latent representations, colored by cell types. **c**, UMAP embeddings of raw enhanced and simulated 3D spatial profiles with annotated mLISI (standard deviation shown in bracket). **d**, Bar plot showing the assessment of single 3D slice generation: Pearson correlation between real and generated data via KNN and STADiffuser, with or without cell type labels. Missing slices are annotated on x-axis ticks. **e**, Bar plot showing the assessment of multiple 3D slice generation, ranging from one to four slices; missing slices are annotated on x-axis ticks. **f**, Heatmap displaying representative gene expression patterns in the generated E16-18h_a_S04 slice with a single slice missing. **g**, Heatmap displaying representative gene expression patterns in the generated E16-18h_a_S05 slice, with slices E16-18h_a_S05-06 missing.

STADiffuser enables the generalization of unseen 3D slices resembling raw data. We trained STADiffuser by excluding one slice and generated that slice both with and without cell-type labels provided and employed K-nearest neighbors (KNN) as a baseline. In this approach, we reconstructed the missing slice using neighboring spots from the adjacent slice. When cell type labels were available, we restricted the selection to neighboring spots of the same cell type. Our results consistently demonstrate the superior performance of STADiffuser over KNN (**Fig. 6d**). Notably, incorporating cell type information boosted the gene expression reconstruction. STADiffuser consistently outperformed the baseline method under both conditions, with and without cell type information. However, cell-type labels for the missing slice are often unavailable in practical scenarios. Interestingly, even without cell type labels, STADiffuser outperformed KNN with cell type labels in most cases (except for the excluded slice E16-18h_a_S10).

We then removed multiple slices from one to four to assess the 3D slice generation capability under more challenging conditions. The performance gap between STADiffuser and KNN becomes clearer (**Fig. 6e**). The Pearson correlation of KNN steadily decreases, highlighting its inherent limitations in reconstructing spots physically distant from the profiled ones. In contrast, STADiffuser maintains a high level of correlation even with four slices removed, especially when provided with cell type labels (**Fig. 6e**, E16-18h_a_S04-07, indicating slice S04-07 is missing). Spatial mappings of three representative genes demonstrate that STADiffuser accurately reconstructed the missing slices (**Fig. 6e**, with slice S04 removed; **Fig. 6f**, with slices S04-06 removed). For example, *lov*, playing a pivotal role in embryonic development^34^, exhibited recovered spatial patterns in the reconstructed slice by STADiffuser. STADiffuser, even without labels, achieved a Pearson correlation coefficient of 0.82, surpassing the performance of KNN with labels, which yielded a correlation coefficient of 0.70 (**Fig. 6e**, first row). When dealing with multiple missing slices, KNN produced low correlations (Pearson correlation *r* is below 0.1) in reconstructing the gene *LvsB*. Conversely, STADiffuser consistently and accurately reconstructed the spatial patterns of gene *LvsB* within the missing slice E16-18h_a_S05. This suggests that STADiffuser effectively captures the spatial information inherent in the ST data.

### STADiffuser enables *de novo* reconstruction of a full-view 3D marmoset cerebellum atlas from coronal slices

Whole-organ 3D reconstruction remains challenging due to two key issues: the computational demands of large-scale data and the limitations of physical sectioning, where inter-slice distances greatly exceed intra-slice resolution, resulting in highly anisotropic inputs. To address these challenges, STADiffuser reconstructs a high-resolution marmoset cerebellum atlas from sparse coronal slices, enabling scalable generation and *de novo* synthesis of virtual slices from arbitrary orientations for comprehensive full-view 3D modeling.

We used 33 coronal slices of the marmoset cerebellum^35^ with an inter-slice distance of 250 μm and an intra-spot resolution of 25 μm. Following preprocessing, the dataset comprised 1,919,455 spots and 2,371 highly variable genes, offering a large-scale input for training STADiffuser. Existing methods are not well equipped to handle such large, sparse, and anisotropically sampled datasets.

As a preparatory step, we used *3D Slicer* to manually align the coronal slices and reconstruct a coarse 3D surface model. We trained STADiffuser on the aligned 3D spatial coordinates (**Figure 7a**). To accommodate the dataset’s scale and increase model capacity, we employed a denoising network with 11.81 million parameters trained on two NVIDIA L40 GPUs for 12 days (see **Supplementary Materials** “**Implementation details**”).

**Fig. 7.**
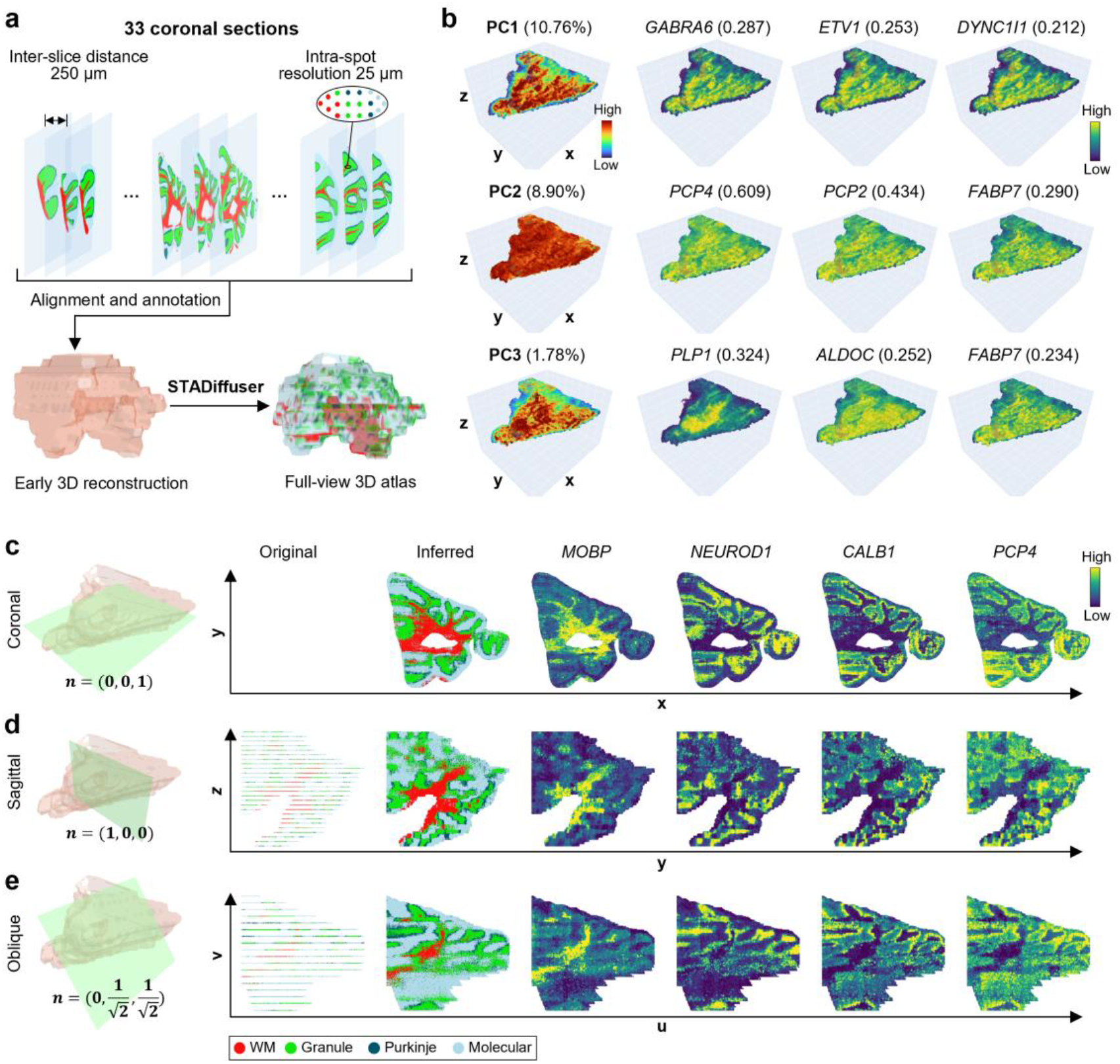
STADiffuser reconstructs a full-view 3D marmoset cerebellum atlas from coronal slices. **a**, Workflow for reconstructing the 3D atlas from 33 coronal sections. **b**, Visualization of the top three sparse principal components (PCs) in 3D, showing explained variance ratios and the top three genes with the highest loading coefficients for each PC. **c-e**, Generated slices in three orientations: coronal **c**, sagittal **d**, and oblique **e**, with the corresponding normal vectors annotated. The original and STADiffuser-generated slices were labeled using a random forest classifier trained on observed spot embeddings; note that no original spot data are available for the coronal slice. Four representative genes highlighting distinct cerebellar layers are shown: *MOBP* (white matter, WM), *NEUROD1* (granule layer), *CALB1* (Purkinje layer), and *PCP4* (molecular layer). For the oblique slice **e**, the projection basis vectors **u** and **v** are defined by the first two PCs of the spot spatial coordinates.

To address the effects of anisotropic sampling, we used STADiffuser to generate a high-resolution 3D atlas comprising 321 virtual coronal slices, reconstructing over 20 million spatial spots and enabling dense, continuous visualization of the cerebellum. The 3D spatial distribution of the top three principal components (PCs) with their highest-loading genes indicates their spatial variations (**Figure 7b**). PC1 primarily corresponds to the granule layer, marked by genes such as *GABRA6* and *ETV1*. PC2 captures variation in the outer layers, including the Purkinje and molecular layers. PC3 is associated with the inner regions, showing strong loadings for white matter marker genes such as *PLP1* and *ALDOC*. These coherent spatial patterns underscore the ability of STADiffuser to recover biologically meaningful 3D structures, enabling integrative analysis across densely reconstructed virtual slices.

Notably, the trained model also enables *de novo* slice reconstruction at arbitrary angles. Given a point on a plane and its normal vector, we compute its intersection with the surface model and sample the corresponding slice using STADiffuser. We show the reconstructed coronal, sagittal, and oblique slices (**Figures 7c–e**). The coronal slice preserves the expected laminar structure marked by known gene-layer associations: *MOBP* (white matter), *NEUROD1* (granule layer), *CALB1* (Purkinje layer), and *PCP4* (outermost layer) (**Figure 7c**). To facilitate automated annotation, we trained a random forest classifier on the latent embeddings. While the classifier broadly captures the overall structure, it may not reliably distinguish finer boundaries, such as those of the Purkinje layer. Despite this limitation, STADiffuser-generated slices, including the more challenging sagittal view (**Figure 7d**), retain coherent laminar organization. The model also supports the reconstruction of oblique slices (**Figure 7e**), which are rarely obtainable in biological experiments. These generated views reveal spatial patterns and gene expression domains that are difficult to observe in conventional sections. In short, STADiffuser scales to millions of spatial locations and enables flexible 3D tissue exploration from arbitrary views, extending beyond the constraints of experimental slicing. An interactive demo showcasing arbitrary-view slice generation is available at https://zhanglab-amss.org/Omni-View-3D-Cerebellar/.

## Discussion

We present a flexible deep generative tool, STADiffuser, to simulate and generate high-fidelity spatial transcriptomic data, serving various downstream tasks (e.g., data imputation, *in silico* experiments, and cell-type-specific gene identification). Furthermore, STADiffuser offers a unified approach for modeling 3D slices and multiple samples, facilitating the utilization of extensive spatial transcriptomic data.

The foundation models, enabling cell typing, universal representation, and various tasks, are emerging in single-cell multi-omics data^36–38^. Yet, foundation models tailored for spatial transcriptomics remain underexplored. STADiffuser, capable of handling multiple samples and scalable to millions of spots, presents a promising avenue for developing foundation models in spatial transcriptomic data. Nevertheless, STADiffuser for multi-sample modeling relies on manually aligning coordinates, which restricts its application to general tissues. It is expected that future work will develop a universal coordination aligning strategy or a unified coordination representation strategy, enabling the training of massive ST data in a powerful generative model. This is a crucial step towards the foundation model in ST data.

STADiffuser currently only models the spatial transcriptomic data. With the development of spatial multi-omics technology, it can also be extended to simulate spatial multi-omics data, deciphering the complex interactions between different molecular layers in tissues. This integration will provide a more comprehensive understanding of spatial organization and functional relationships within complex biological systems.

## Materials and Methods

### Data preprocessing

STADiffuser processes gene expression profiles and spatial coordinates of single or multiple slices as its input. We conducted normalization and log transformation for the raw gene expressions using the SCANPY^39^ package. We identified highly variable genes through the `*scanpy*.*pp*.*highly_variable_genes*` function. To account for scale differences across spatial transcriptomics technologies, we rescaled spatial coordinates so that the average inter-spot distance is standardized, ensuring compatibility with spatial positional encoding mechanisms.

### Constructing the spatial neighbor network

We constructed a spatial network based on Euclidean distance to model local spatial information. Specifically, edges were assigned between spots if their distances were within a predefined threshold distance, denoted as *r*. This spatial neighbor network (SNN) was constructed individually within each slice. The *r* was chosen such that the average number of neighbors was approximately 5. For single-nuclei resolution technologies, such as Slide-tags, we chose not to utilize SNN due to its already high resolution.

### Deep graph attention autoencoder

To learn the representation of spots, we designed a deep graph attention autoencoder integrated with several modern neural network techniques. Specifically, we used two graph attention layers to adaptively aggregate information of neighbor spots and reduce dimensions due to its effectiveness in spatial transcriptomic data modeling, like STAGATE^7^. In subsequent, a self-attention block^40^ was adopted to further process the data for its excelling at complex data distribution.

#### Graph attention layer

Given the normalized gene expression of the *i*-th spot, denoted by ***x***_*i*_ ∈ *R*^*p*^, where *p* is the number of genes, the representation of the *i*-th spot 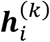 graph attention layer at the *k*-th layer follows:

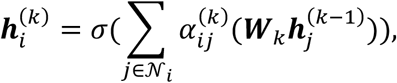

where 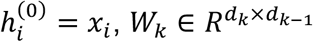 is a learnable weight matrix in the *k*-th layer, 𝒩*i* is the neighbor set of spot 𝒩*i* in SNN, *α*_*ij*_ is the learnable attention weight between the two spots, and *σ* is a nonlinear activation function.

#### Self-attention block

The output of the graph attention layers first transforms by a linear projection:

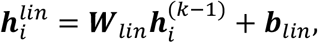

where *b*_1_ is a learnable intercept. Then a single-head attention layer follows:

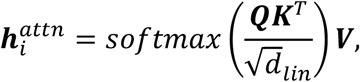

where 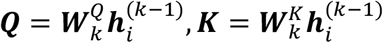, and 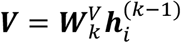 indicate query, key and value matrices, respectively, and 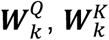, and 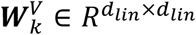 are corresponding weight matrices. The output of the attention layer 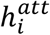 attended with the input was passed to a two-layer multilayer perceptron (MLP), and the final output of the self-attention block 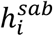 was computed as the following:

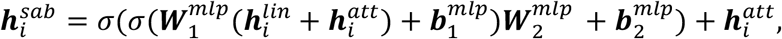

where 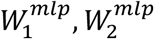 and 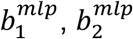 indicate the weight matrices and intercept vectors in the MLP, respectively. Layer normalization^41^ is applied to 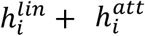 and 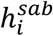 to stabilize the training.

#### Overall architecture

The encoder *f*_*enc*_ is composed of two graph attention layers (512-32), which aggregate information and reduce the dimensionality to 32, followed by a self-attention block that further processes the representation to output a 32-dimensional spot representation. Denoting the *i*-th spot representation as *z*_*i*_ ∈ *R*^*d*^, a decoder *f*_*dec*_ of the symmetric architecture of encoder reconstructs *z*_*i*_ back to the gene expression space 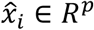. ADAM^42^ optimizer is employed to minimize the reconstruction loss:

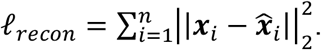

To simplify notation, we’ll omit the subscript *i* when it’s clear from context.

### Triplet learning for multi-sample graph attention autoencoder

We adopted a triplet loss technique to mitigate batch effects across multiple sections/slices/samples. This method has proven effective in integrative spatial transcriptomic data analysis, e.g., STAligner^11^. Specifically, triplet learning leverages anchor, positive, and negative samples to ensure that the learned representation of each anchor is closer to its positive counterpart than to the negative one. Like STAligner, we defined the anchor and positive pairs as the mutual nearest neighbors (MNN) across two sections and the anchor. Notably, the MNN pairs were computed in the latent space *z* and updated during the training progress, to avoid potential noise and for further refining.

The negative points were randomly sampled from the same section. The triplet loss is given as the following:

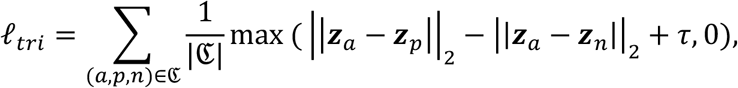

where 𝔊 indicates the triplet set indices of, (*a, p, n*) denotes the indices of anchor, positive and negative points, respectively, and *τ* > 0 is a margin between positive and negative pairs (default value 1). The autoencoder is first pretrained on each slice with reconstruction loss for *T*_1_ epochs and then optimized by the combing loss:

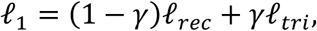

where *γ* > 0 is a hyperparameter with a default value of 0.5. The autoencoder is trained with *T*_2_ epochs by updating the triplet indices set 𝔊 every *m* epochs. In the experiments, we set *T*_1_ = 200, *T*_2_ = 300 and *m* = 50.

### Spatial positional encoding

To capture the long-range spatial information, we adopted sinusoidal positional encoding to encode the spot coordinates, that were originally used to represent the location or position in a sequence in transformer^40^. We extended it to inform the diffusion model of the 2D/3D spatial coordinates. The idea is straightforward. We start with 2D coordinates. Let’s denote the quantized spatial coordinates of spot *i* as 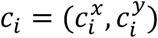, the x-axis positional embedding is a 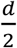-dimensional vector 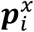, whose 2*j*-th or (2*j*+1)-th element is:

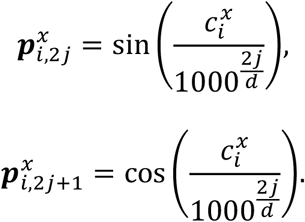

The y-axis positional embedding 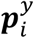 is defined in the same way. Hence, the 2D space positional embedding is given by: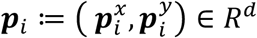.

For 3D coordinates, since the z-axis, i.e., the thick of the section, is typically not of the same scale as the x and y-axis, we defined the z-axis positional embedding as a *d*-dimensional vector 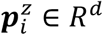 with the same procedure as 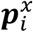.Finally, the 3D space positional embedding of the *i*-th spot is given by concatenating the 2D space positional embedding with the z-axis embedding, i.e., 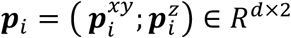,where 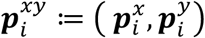.

### Latent space diffusion model

The key idea of the diffusion model is to learn a denoise network that gradually transforms the prior distribution (standard Gaussians) to the data distribution. The gene expression profile of ST data is both highly sparse and high-dimensional, making direct modeling and sampling challenging. Hence, we adopted a diffusion model in the latent space to enable sampling from the spot latent embedding space and further reverse back to the gene expression using the trained decoder.

The diffusion model consists of a forward process that gradually adds noise to the data and a backward process that gradually denoises the noise to the data through a spatial denoising network.

#### The forward process

Specifically, the forward process is a Markov chain with a variance schedule 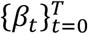:

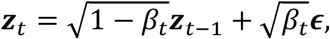

where *t* = 1, …, *T, z*_0_ = *z* is the latent embedding of a spot, *ϵ* ∈ 𝒩 (**0, *I***_***d***_) is the standard Gaussian adding noise, and 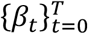 is a schedule variance that controls the scale of the adding noise with *β*_*t*_ ∈ (0,1). The *z*_*t*_ at arbitrary timestep *t* admits the close-form equation:

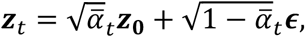

where *α*_*t*_ ≔ 1 − *β*_*t*_ and 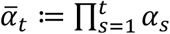. It is evident that when 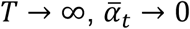, and thus *z*_*T*_ → *ϵ*. Hence, the noisy data *z*_*T*_ can be regarded as following a standard normal distribution with sufficient larger *T* and appropriate schedule variance.

#### The backward process

The backward process reverses the noise ***z***_*T*_ ~ 𝒩 (**0, *I***_***d***_) gradually by a spatial denoising network *ϵ*_***θ***_ parametrized by ***θ***:

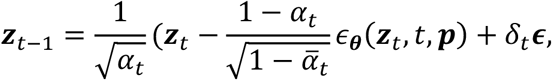

where *δ*_*t*_ = 0 if *t*=1, otherwise *δ*_*t*_ = 1. *ϵ*_***θ***_ is the spatial denoising network that takes the noisy data ***z***_*t*_, current timestep *t* and positional embedding ***p*** to predict the noisy data *z*_*t*−1_ at *t* − 1 timestep. *z*_0_ is expected to follow the spot latent distribution and can be further reversed back to the gene expression space by the decoder. The spatial denoising network is trained to minimize the reconstruction error of noise via the loss function defined as:

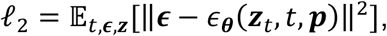

where the expectation is empirically approximated by randomly drawing samples from *t* ∈ {1, · ·, *T*}, ***ϵ*** ∈ 𝒩 (**0, *I***_***d***_) and ***z***_***t***_ from the noisy spot representations.

### Spatial denoising network

The spatial denoising network *ϵ*_***θ***_ adopts a U-Net backbone^43^ with enhanced self-attention layers. The spatial positional embedding ***p*** is fed to *ϵ*_***θ***_ to preserve the spatial patterns. Specifically, in the 2D scenario, the spatial positional embedding ***p*** is added to the timestep embedding, denoted as ***e***_*t*_:

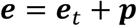

where the timestep *t* is projected to an embedding vector ***e***_*t*_ ∈ *R*^*d*^.

When the conditional labels, denoted as *l*, are available, we enhance our framework with a conditional spatial denoising network. This enables the synthesis of data under specified conditions. In this case, the embedding vector is extended to incorporate the conditional information:

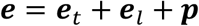

where label *l* is transformed into the condition-specific embedding ***e***_*l*_ ∈ *R*^*d*^. Both the embedding vector ***e*** and ***z***_*t*_ are used to predict the noise ***ϵ***. For 3D data, the embedding vector is computed as:

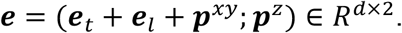

This composite embedding vector ***e*** undergoes an MLP transformation tailored for the appropriate dimensions before being processed by the U-Net backbone network.

### Evaluation metrics of simulation methods

We evaluated simulation methods based on fidelity, measuring how closely simulated data mirrors the truth, and diversity, gauging the variability within simulated data.

#### Fidelity

Given the reference gene expression matrix ***X*** ∈ ℝ^*n*×*p*^ of *n* spots and *p* genes, and the corresponding simulated data 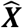, the fidelity is defined as the following:

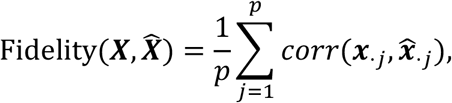

where 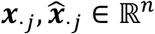 are the profiles of *j*-th gene in ***X*** and 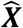, respectively. *corr* is the Pearson correlation *r*.

#### Diversity

Given *M* simulated replicates 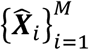, the diversity is defined as the following:

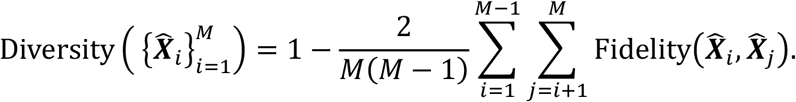

The diversity is defined as the one minus the average of the pairwise fidelity across simulated replicates. We set *M*=5 in all experiments.

### Computing the co-localization metric

We defined a co-localization metric (co-loc) to quantify the spatial consistency between two subsets of spots. Given two indices sets, a reference set 𝒥 and a target set 𝒥, the co-loc is defined as:

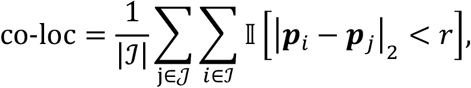

where 𝕀 is an indicator function, ***p***_*i*_ and ***p***_*j*_ are the spatial position of the *i*-th and *j*-th spots, respectively. *r* is a distance thresholding. We typically set *r* as the average of the nearest neighbor’s distance. To assess the spatial consistency between a gene and a reference set 𝒥, we selected the top | 𝒥 | highly expressed spots as the target set.

### Identifying cell-type-specific genes

We trained a STADiffuser model with spot gene expression profile 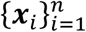 and corresponding cell type labels 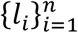, where *n* is the number of trained spots. To identify cell-type-specific genes between cell types *c*_1_ and *c*_2_ with controlling the confounding spatial position, we simulated an *in silico* replicate 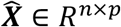 and a pseudo *in silico* replicate 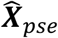 by swapping the cell type assignments of *c*_1_ and *c*_2_ in 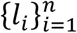. The mean expression differences were calculated for all genes across the simulated datasets of cell types *c*_1_ and *c*_2_:

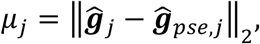

where 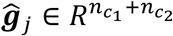 is the *j*-th gene profile vector of cell types *c*_1_ and *c*_2_ spots in 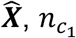 and 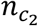 denote the numbers of spots for each cell type, and ĝ_*pse,j*_ is the corresponding 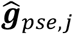 gene vector in 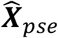.Finally, the cell-type-specific gene is selected by the following criterion:

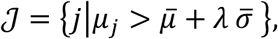

where 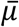 and 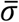 represent the mean and standard deviation across all genes, calculated as 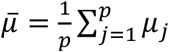 and 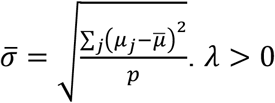 is a user-defined parameter to control the number of selected cell-type-specific genes. We set *λ* = 1·5 in the experiments.

### Differentially expressed genes, gene enrichment analysis, and gene scores

We identified differentially expressed genes by comparing target cells/spots to a reference using the Wilcoxon rank-sum test. We used the g:pforfier API^44^ provided by SCANPY^39^ to perform gene enrichment analysis. We scored cells/spots by applying function *tl*.*score_genes* in SCANPY with a given marker gene list.

## 3D surface reconstruction

We first voxelized the 3D spatial coordinates from all 33 coronal slices using Gaussian smoothing and converted the result into an NRRD (Nearly Raw Raster Data) file, which supports multi-dimensional spatial data and is compatible with medical imaging tools. Using the graphical interface software *3D Slicer*^45^, we manually aligned the slices to correct for inter-slice misalignment. We then annotated anatomically hollow regions within the cerebellum, such as internal fissures and cavities, to ensure their preservation when reconstruction. Contour-based interpolation was applied along the anterior-posterior (z) axis to generate a coarse 3D cerebellum model that maintains these internal voids. The resulting 3D surface model provides a geometric reference that enables the computation of intersections with arbitrary slicing planes for virtual visualization.

### Dimensionality reduction and automated layer annotation

We employed `*MiniBatchSparsePCA*`, as implemented in *scikit-learn*^46^, to perform sparse principal component analysis on large datasets while avoiding memory constraints. To this end, we used a mini-batch size of 100 and retained the default *ℓ*_1_ penalty parameter of 1.0 to enforce sparsity in the loading vectors.

To approximate the explained variance ratio (EVR) for each sparse principal component (PC), we computed the variance of the projected data and normalized it by the total variance of the centered input data. Specifically, let *X*_*centered*_ ∈ *R*^*n*×*p*^ be the mean-centered data matrix, and PC_k_ = *X*_*centered*_ ***w***_*k*_ be the projection onto the *k*-th component. The EVR is approximated as:

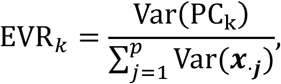

where Var(·) indicates the variance.

To automate layer annotation, we trained a random forest classifier with 100 trees using the latent embeddings of the observed spots as input features. The dataset was split into training and test sets at a 4:1 ratio. The classifier achieved an overall test accuracy of 0.86. However, performance on the Purkinje cell layer was lower, with a precision of 0.68 and recall of 0.65, likely due to its unique structure as a narrow band situated between the molecular and granule layers, which makes it more difficult to resolve and annotate accurately. Detailed accuracy metrics are provided in **Table S3**.

## Data availability

All data used in this paper are available in raw from the corresponding studies. Specifically, the DLPFC dataset generated by 10x Visium is available at http://spatial.libd.org/spatialLIBD. The hippocampus dataset of the J20 mouse model generated by Slide-seq V2 is accessible at Broad Institute’s Single Cell Portal with accession number SCP1663. The validated J20 mouse model scRNA-seq hippocampus dataset is available at NCBI with accession number GSM4783652. The human melanoma dataset and multiome dataset generated by Slide-tags are available at Broad Institute’s Single Cell Portal with accession numbers SCP2171 and SCP2176, respectively. The 3D datasets of *Drosophila* generated by Stereo-seq are available at https://db.cngb.org/stomics/flysta3d/. The coronal slices of the marmoset cerebellum dataset are available at https://db.cngb.org/stomics/cbmsta/.

## Acknowledgments

This work has been supported by the Strategic Priority Research Program of the Chinese Academy of Sciences (no. XDA0460303 to C.Z.), the National Key Research and Development Program of China (nos. 2025YFF1207900 to S.Z. and 2024YFF0729201to C.Z.), the National Natural Science Foundation of China (nos. 32341013, 12326614 to S.Z. and 12401661to C.Z.), The Zhejiang Province Vanguard Goose-Leading Initiative (no. 2025C01114), and the CAS Project for Young Scientists in Basic Research (no. YSBR-034 to S.Z.).

## Author contributions

Shihua Zhang conceived and supervised the project. Chihao Zhang designed and implemented the STADiffuser algorithm. Chihao Zhang and Shihua Zhang validated the study. Chihao Zhang and Shihua Zhang wrote the manuscript.

All authors read and approved the final manuscript.

