## Supplemental Text and Figures for "STADiffuser: high-fidelity simulation and full-view 3D modeling of spatial transcriptomics"

### Supplementary Notes

#### Datasets

We applied STADiffuser to various datasets generated by different ST platforms to illustrate its effectiveness in this study. Specifically, we applied STADiffuser to the human dorsolateral prefrontal cortex (DLPFC) dataset[1], J20 mouse model of Alzheimer's disease generated by Slide-seq V2[2], mouse visual cortex dataset generated by STARMap[3], human melanoma tumor dataset obtained through Slide-tags profiling[4], *Drosophila* E16-18 embryo dataset generated by Stereo-seq[5], and marmoset cerebellum dataset consisting of coronal slices generated by Stereo-seq[6]. The details of these datasets are shown in **Table S1**.

#### Implementation details

The autoencoder takes an input with a variable number of genes. It passes them through graph attention layers with hidden dimensions of 512 and 32, followed by attention blocks output dimensions of 32 and 32. The model uses the ELU activation and dropout for regularization.

The denoising network takes the form of a 1D U-Net. For conventional datasets, we employ the smaller architecture with moderate depth and channel size to balance performance and efficiency. For the marmoset cerebellum dataset, which contains millions of spots, we use the larger model with increased depth and channel capacity to capture complex spatial patterns and ensure sufficient representational power. Please refer to **Table S2** for a detailed comparison of the architectures and their parameter counts.

#### STADiffuser training specifications

We trained the graph autoencoder using the mini-batch mode with the ADAM optimizer[7] and a learning rate of 1e-4. The batches were constructed using neighbor sampling[8], with the number of neighbors set to [5, 3], indicating the numbers of first-order and second-order neighbors. For multiple slices, we pre-trained the autoencoder on each slice for 200 epochs. We then further trained the autoencoder with a triplet loss constructed by mutual nearest neighbors (MNNs) in the latent space for an additional 300 epochs. The MNNs were updated every 50 epochs.

The latent space diffusion models were also trained using the mini-batch mode using the ADAM optimizer with a small learning rate of 1e-5. The denoising networks were trained using the same sampling procedure for a total of 1000 epochs. For conventional datasets, the training takes several hours on a single NVIDIA L40 GPU. In contrast, training on the cerebellum dataset updated for 800 epochs costs 12 days using 2 NVIDIA L40 GPUs.

#### Comparison with competing methods

We compared STADiffuser with five commonly used methods for transcriptomics data simulation: Splatter and its kernel variant (KerSplatter)[9], ZINBWaVE[10], scDesign3[11], and SRTsim[12]. Specifically, Spatter and KerSpatter are statistical frameworks based on the gamma-position distribution. They are specifically designed for single-cell RNA-seq (scRNA-seq) datasets without accounting for spatial information. ZINBWaVE is a flexible statistical approach to modeling the count data with a zero-inflated negative binomial model. It can incorporate the spatial information as covariates. scDesign3, a recently developed versatile statistical simulator, can incorporate spatial information, cell type, and time trajectories as covariates. In particular, it introduces copula models to capture the gene-wise correlation to enable high-fidelity simulation. However, its computational cost is expensive.

We note that all the compared methods are directly modeling the count data. Hence, the count data is normalized by `*pp.normalize_total*` with parameter `*target_sum=1e4*` in SCANPY package[13]. To facilitate a fair comparison, the simulated datasets were enhanced by the graph autoencoder trained on the real data. Then the evaluation metrics, including the gene-wise Pearson correlation *r*, the mean of local inverse Simpison's index (mLISI), were computed. We followed the authors' instructions for detailed implementation of the compared methods. The spatial coordinates were normalized to [0,1] with the min-max normalization.

- **Splatter and KerSplatter:** we used the functions `*splatSimulate*` and `*kersplatSimulate*` with default parameters. The coordinate information was not used.
- **ZINBWaVE:** we used the `zinbEstimate` function with the parameter `design.sample` setting as normalized spatial coordinates to estimate the model parameters. Then the simulated data were sampled by `*zinbSimulate*` with the estimated parameters.
- **scDesign3**: we used the wrapper function `*scdesign3*` with normalized coordinates and either cell type or spatial domain labels to estimate model parameters to estimate the model parameters. Specifically, the model used negative binominal to model the count data. We set the number of Gaussian copula components to 20.
- **SRTSim**: we used ` *srtsim_fit*` with `*sim_scheme=domain*` and normalized spatial coordinates to fit the model parameters. Cell type or spatial domain labels were used when available. The simulated data were generated by `*srtsim_count*` with default parameters.

**Comparative analysis of STADiffuser and SRTsim in super-resolution**

We compared STADiffuser and SRTsim in terms of super-resolution on the DLPFC dataset, as other competing methods do not offer a direct solution for super-resolution. We randomly removed spots to create the training data, with missing rates ranging from 0.1 to 0.5. As shown in **Fig. S1a**, STADiffuser consistently outperformed SRTsim. The performance of both methods decreased as the missing rates increased. Notably, the performance gap between STADiffuser and SRTsim narrowed with higher missing rates, suggesting that the deep learning approach of STADiffuser is more promising when the dataset is sufficiently large.

We also showed three representative genes, *GAD1*, *KRT17*, and *B3GALT2,* for detailed comparison (**Fig. S1b-d**). STADiffuser accurately reconstructed the original data (upper-left and first-row panels) from the downsampled data (third-row panels).

### Supplementary Figures

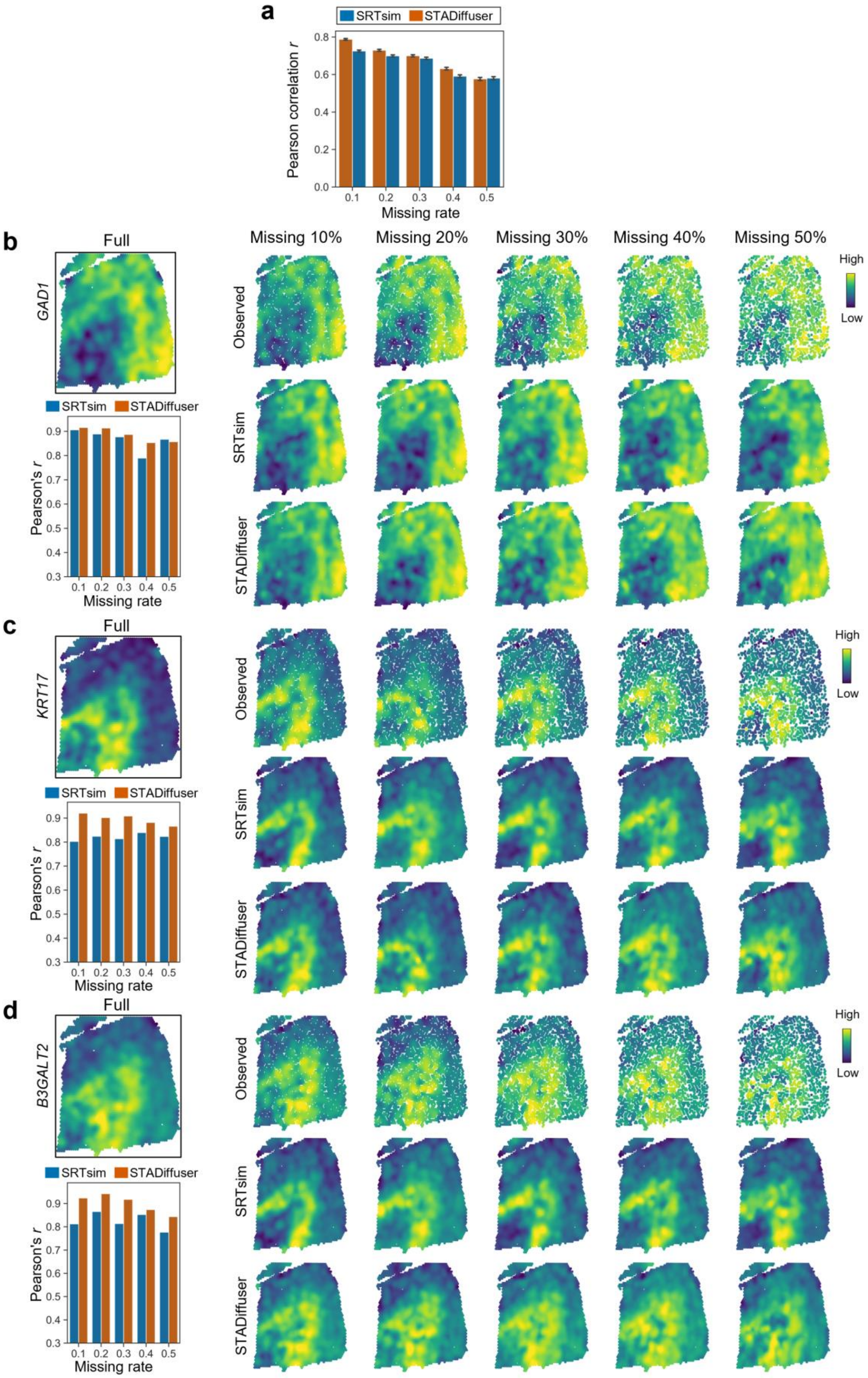

**Fig. S1 Comparison of super-resolution performance on the DLPFC dataset**. **a,** The average of the gene-wise Pearson correlation concerning the missing rates. **b-d,** Comparison for the super-resolution performance of STADiffuser and SRTsim in three representative genes, i.e., *GAD1*, *KRT17*, *B3GALT2*. The upper-left panels show the spatial mapping of gene expression, and bar plots in the bottom-left panels show the Pearson correlations of the recovered data. The three rows on the right present, from top to bottom: observed data, gene expression recovered by SRTsim, and gene expression recovered by STADiffuser.

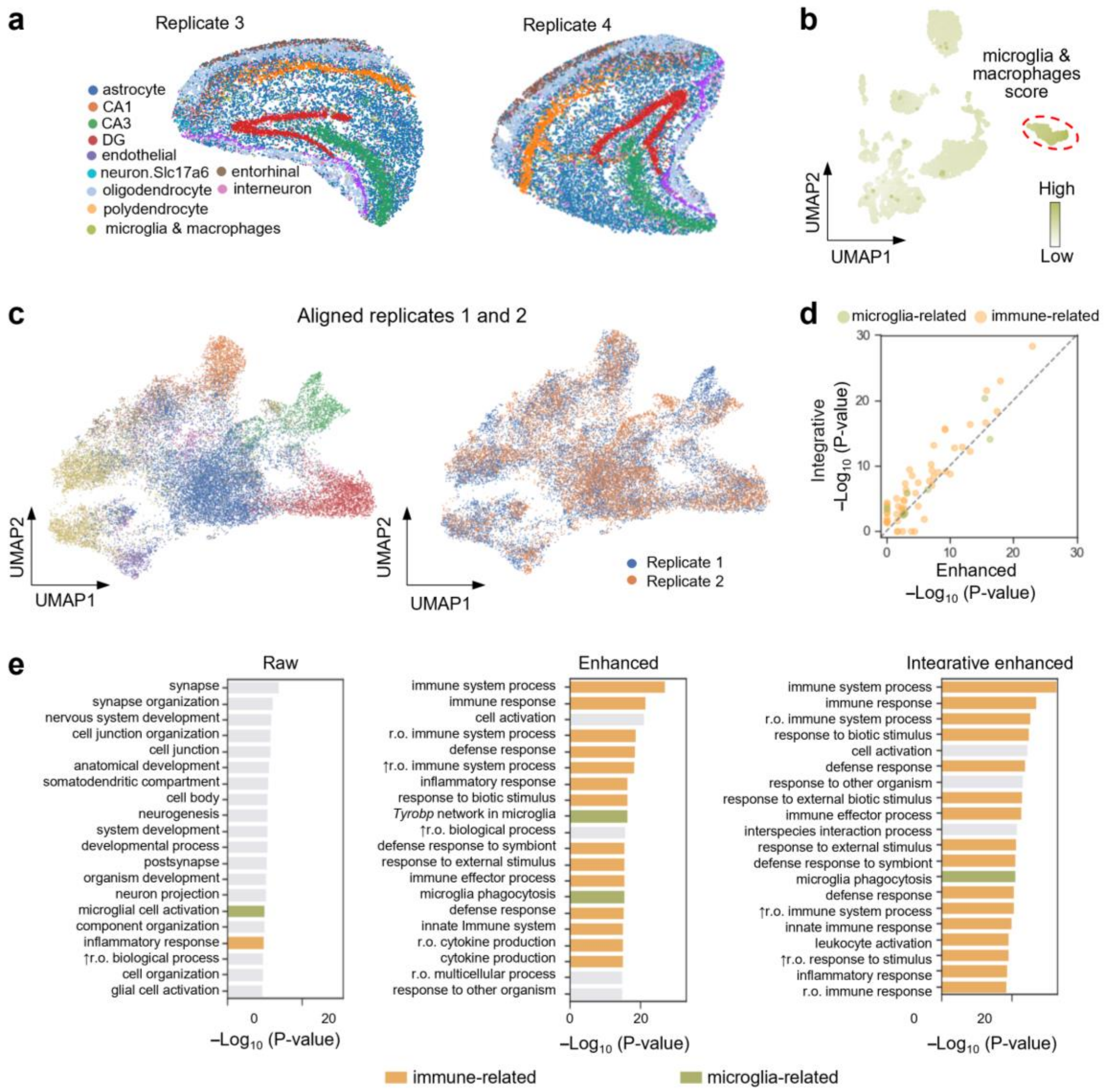


**Fig. S2 Additional results on the J20 mouse hippocampus dataset**. **a**, Spatial mappings of replicates 3 and 4. **b**, UMAP embeddings of the scRNA-seq reference data, colored by microglia and macrophage scores. The identified microglia&macrophages cluster is highlighted with a red circle. **c,** UMAP embeddings of the aligned representations of replicates 1 and 2, colored by cell types (same color scheme as in panel **a**). **d,** Scatter plot showing the log-transformed GO enrichment terms of enhanced data (single slice) versus integratively enhanced data (multiple slices). **e,** Bar plots showing the top enriched GO terms in raw, enhanced, and integratively enhanced data. Immune-related and microglia-related enriched GO terms are colored orange and green, respectively; non-specific GO terms are shown in grey.

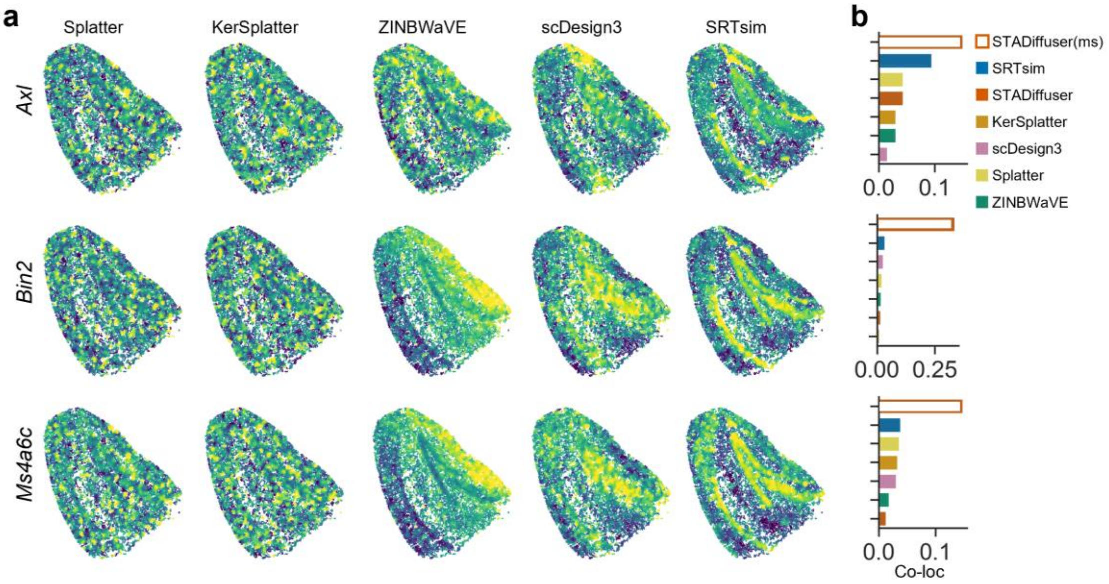


c

| Gene | $-\log_{10}(P_{adj})$ | Gene | $-\log_{10}(P_{adj})$ | Gene | $-\log_{10}(P_{adj})$ |
|---|---|---|---|---|---|
| *Adgre1* | 125.0587 | *Igf1* | 37.2180 | *Prcp* | 22.6660 |
| *Axl* | 17.9440 | *Il10ra* | 125.0587 | *Ptpn18* | 119.5729 |
| *Bin2* | 125.0587 | *Itgb2* | 124.0704 | *Rin2* | 89.2557 |
| *Ccl9* | 124.6143 | *Jam2* | 73.9392 | *Slc14a1* | 72.9504 |
| *Cd63* | -0.5395 | *Lcp2* | 125.0587 | *Slc40a1* | 124.9048 |
| *Cspg4* | 96.8706 | *Lima1* | 38.9302 | *St3gal6* | 123.6746 |
| *Fermt3* | 125.0587 | *Ltc4s* | 120.1611 | *Tgfb1* | 125.0587 |
| *Fgd2* | 125.0587 | *Mob1a* | 25.8430 | *Tnfaip8l2* | 124.6187 |
| *Gpr183* | 125.0587 | *Ms4a6c* | 124.7519 | *Ucp2* | 115.4722 |
| *Higd1b* | 85.5323 | *Plcb3* | 29.6557 | *Vamp8* | 99.8404 |

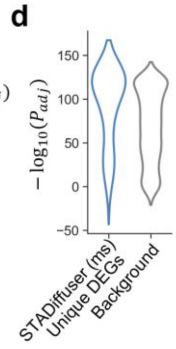


**Fig. S3 Comparative analysis of gene expression and differentially expressed genes (DEGs) using STADiffuser and other methods on the J20 mouse hippocampus dataset. a**, Simulated spatial gene expression profiles generated by five compared methods. **b**, Bar plots showing the co-localization (co-loc) scores for STADiffuser with single and multiple slices (denoted as 'ms'), along with the five compared methods. **c**, DEGs uniquely identified by STADiffuser using multiple slices but not by the single-slice model or SRTsim. The negative log-transformed adjusted p-values of the Wilcoxon rank-sum test in the scRNA-seq reference data are colored by quantiles. **d**, Violin plot displaying the distribution of the negative log-transformed adjusted p-values of the DEGs in Table (**c**) and the rest of the genes.

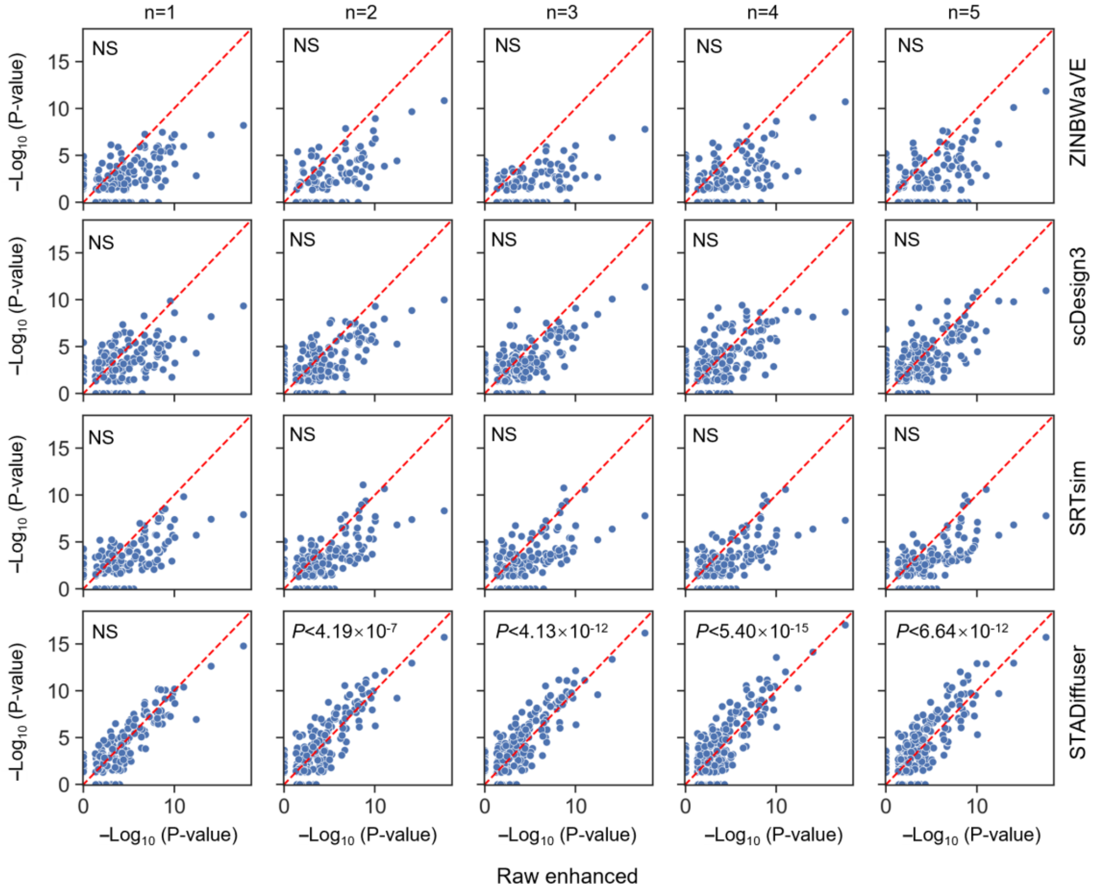


**Fig. S4 Comparative analysis of enrichment significance of GO terms on STARmap mouse visual cortex dataset.** Comparison of GO term enrichment significance based on DEGs from raw enhanced data, STADiffuser, and three other methods, across varying numbers of in silico replicates (from 1 to 5). Dotted diagonal lines depict y=x. Wilcoxon rank sum tests were performed on the $-\log_{10}$ (*P* values) between the raw enhanced data and the simulated data. *P*-values are annotated at the upper left in the panels. 'NS' indicates no significance.

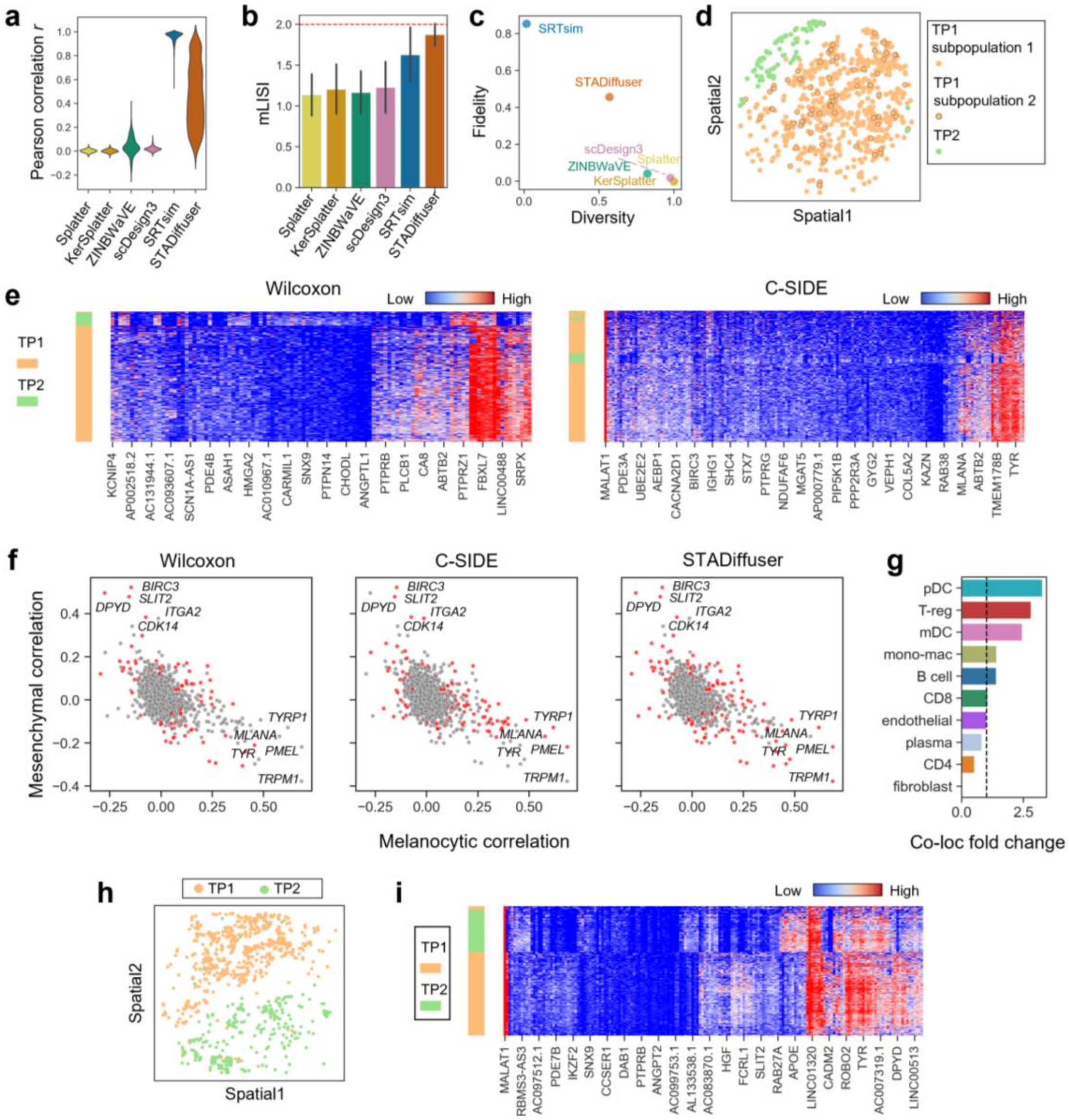

**Fig. S5 Additional results on the Slide-tags human melanoma tumor dataset. a**, Violin plot showing Pearson correlation between raw enhanced data and data generated by the simulation methods. **b**, Bar plot showing the mLISI of the simulation methods, with the optimal mLISI indicated by a horizontal dashed line. **c**, Scatter plot illustrating the characteristics of simulation methods in terms of fidelity and diversity. **d**, Spatial mappings of TP1 subpopulations 1 and 2, along with TP2. **e**, Heatmap illustrating the identified DEG profiles by Wilcoxon rank sum test and C-SIDE, respectively. **f**, Scatter plots of the gene correlations with melanocytic and mesenchymal scores, with genes identified by the methods highlighted in red. **g**, old changes in co-localization values of cell types in TP1 subpopulation 2 versus subpopulation 1; the dashed line indicates a fold change of 1. **h**, Spatial mappings of TP1 and TP2 on the adjacent slice. **i**, Heatmap of gene profiles from the section in panel **h**, clustered by cell-type-specific genes identified from the section in panel **d** by STADiffuser.

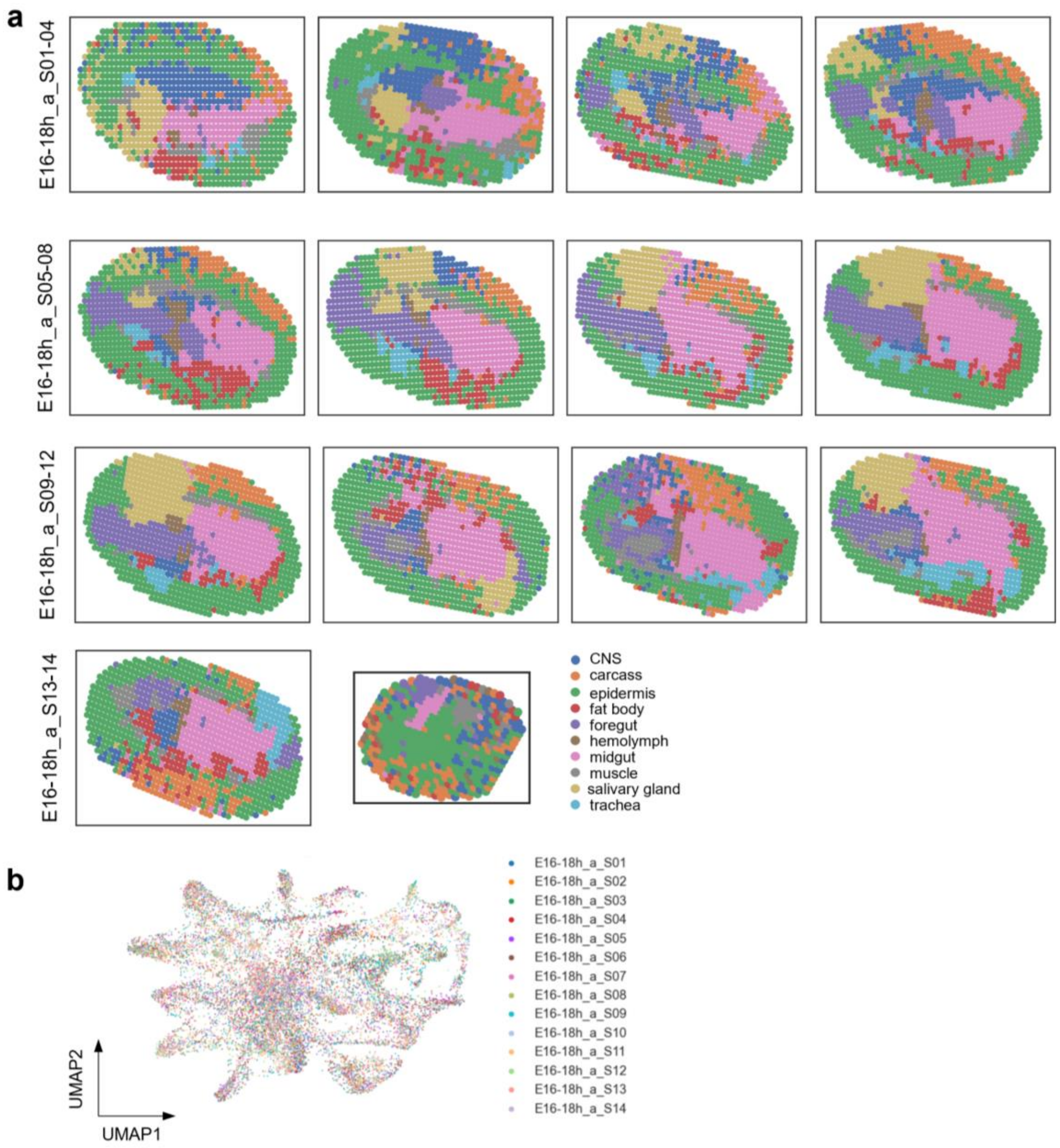


**Fig. S6 Additional results of the Stereo-seq *Drosophila* embryo dataset. a**, Spatial mappings of the 14 slices, colored using a consistent color scheme. Slice E16-18h_a_S14 is smaller than the other slices because it represents the posterior end of the *Drosophila* embryo, which is narrower and less voluminous. **b**, Aligned UMAP embeddings colored by slice ID.

### Supplementary Tables

**Table S1.** Description of all ST datasets used in this study.

| Platform | Tissue | Section | #Spots | Reference |
|---|---|---|---|---|
| **10x Visium** | Human dorsolateral prefrontal cortex (DLPFC) | 151507, 151508, 151509, 151510, 151669, 151670, 151671, 151672, 151673, 151674, 151675, 151676 | 4226, 4384, 4789, 4634 3661, 3498, 4110, 4015, 3639, 3673, 3592, 3460 | [1] |
| **Slide-seq V2** | Hippocampus of J20 genetic mouse model with Alzheimer's disease | Replicate1, Replicate2, Replicate3, Replicate4, | 12918, 10998, 15397, 16800 | [2] |
| **STARMap** | Mouse visual cortex | - | 1207 | [3] |
| **Slide-tags** | Human melanoma tumors | Transcriptomics, Multiomics | 6466 | [4] |
| **Stereo-seq** | *Drosophila* embryo E16-18 | S01, S02, S03, S04, S05, S06, S07, S08, S09 S10, S11, S12, S13, S14 | 985, 965, 1021, 1189, 1181, 1076, 1096 1131, 1113, 1111, 1193, 1049, 1022, 502 | [5] |
| **Stereo-seq** | Marmoset cerebellum dataset | Marmoset1_T478, Marmoset1_T479, Marmoset1_T480, Marmoset1_T482, Marmoset1_T483, Marmoset1_T484, Marmoset1_T485, Marmoset1_T486, Marmoset1_T487, Marmoset1_T488, Marmoset1_T489, Marmoset1_T490, Marmoset1_T491, Marmoset1_T493, Marmoset1_T495, Marmoset1_T496, Marmoset1_T497, Marmoset1_T499, Marmoset1_T501, Marmoset1_T502, Marmoset1_T503, Marmoset1_T504, Marmoset1_T505, Marmoset1_T506, Marmoset1_T509, Marmoset1_T511, Marmoset1_T513, Marmoset1_T514, Marmoset1_T515, Marmoset1_T516, Marmoset1_T517, Marmoset1_T518, Marmoset1_T519 | 2877, 4840, 6944, 16916, 23516, 27390, 38261, 45591, 56880, 62086, 69260, 71672, 70850, 75334, 84895, 87522, 92996, 104576, 100039, 97791, 99983, 99844, 97298, 96917, 80336, 69769, 55041, 44784, 33597, 29055, 27500, 23688, 21407 | [6] |

**Table S2. 1D U-Net denoising network architectures and parameters.**

| Model | Input Size | Architecture (Down → Up) | Channels | Parameters |
|---|---|---|---|---|
| **1D U-Net** | 1×32 | NoSkip → Std → Attn → Attn → Std → NoSkip | 32 → 32 → 64 | ~711K |
| **1D U-Net (large)** | 1×64 | NoSkip → Attn → Attn → Attn → Attn → Attn → Attn → NoSkip | 64 → 128 → 128 → 256 | ~11.81M |

**Note**: Each model is a 1D U-Net architecture, where **"NoSkip"** indicates blocks without skip connections, **"Std"** refers to standard convolutional blocks, and **"Attn"** denotes blocks with attention. Architectures are symmetric unless otherwise noted. Input size refers to the length of the 1D signal per channel. Parameters are estimated under default configurations.

**Table S3. Random forest classification report for cerebellar layer annotation.**

| Class | Precision | Recall | F1-score | Support |
|---|---|---|---|---|
| Granular layer | 0.91 | 0.89 | 0.90 | 113,206 |
| Molecular layer | 0.87 | 0.92 | 0.90 | 168,368 |
| Purkinje layer | 0.68 | 0.65 | 0.67 | 42,455 |
| White matter | 0.88 | 0.79 | 0.83 | 59,287 |
| Accuracy | | | 0.86 | 383,316 |
| Macro average | 0.83 | 0.81 | 0.82 | 383,316 |
| Weighted average | 0.86 | 0.86 | 0.86 | 383,316 |
